# DCyFIR: a high-throughput CRISPR platform for multiplexed G protein-coupled receptor profiling and ligand discovery

**DOI:** 10.1101/2020.01.16.909325

**Authors:** NJ Kapolka, GJ Taghon, JB Rowe, WM Morgan, JF Enten, DG Isom

**Affiliations:** Department of Molecular and Cellular Pharmacology, University of Miami Miller School of Medicine, Miami, FL, 33136 USA; Flow Cytometry Shared Resource, University of Miami Sylvester Comprehensive Cancer Center, Miami, FL, 33136 USA; University of Miami Sylvester Comprehensive Cancer Center, Miami, FL, 33136 USA; Center For Computational Sciences, University of Miami, Coral Gables, FL, 33146-2930 USA

**Keywords:** G protein-coupled receptors, DCyFIR, multiplex, dark GPCR, yeast

## Abstract

More than 800 G protein-coupled receptors (GPCRs) comprise the largest class of membrane receptors in humans. While there is ample biological understanding and many approved drugs for prototypic GPCRs, most GPCRs still lack well-defined biological ligands and drugs. Here, we report our efforts to tap the potential of understudied GPCRs by developing yeast-based technologies for high-throughput CRISPR engineering and GPCR ligand discovery. We refer to these technologies collectively as Dynamic Cyan induction by Functional Integrated Receptors: DCyFIR. A major advantage of DCyFIR is that GPCRs and other assay components are CRISPR-integrated directly into the yeast genome, making it possible to decode ligand specificity by profiling mixtures of GPCR-barcoded yeast strains in a single tube. To demonstrate the capabilities of DCyFIR, we engineered a yeast strain library of 30 human GPCRs and their 300 possible GPCR-Gα coupling combinations. Profiling of these 300 strains, using parallel (DCyFIRscreen) and multiplex (DCyFIRplex) DCyFIR modes, recapitulated known GPCR agonism with 100% accuracy, and identified unexpected interactions for the receptors ADRA2B, HCAR3, MTNR1A, S1PR1, and S1PR2. To demonstrate DCyFIR scalability, we profiled a library of 320 human metabolites and observed new GPCR-ligand interactions with amino acid, lipid, sugar, and steroid metabolites. Remarkably, many of these findings pertained to understudied “pharmacologically dark” receptors GPR4, GPR65, GPR68, and HCAR3. For example, we found that kynurenic acid activated HCAR3 with a nearly 20-fold lower EC_50_ than GPR35, its known receptor. Taken together, these findings demonstrate the power of DCyFIR for identifying novel ligand interactions with prototypic and understudied GPCRs.

**Significance Statement:** G protein-coupled receptors (GPCRs) are the largest class of membrane receptors in humans. As such, GPCR signaling is central to human biology and medicine. While more than 30% of approved drugs target roughly 150 GPCRs, most receptors lack well-defined endogenous ligands and are currently not druggable. To address this challenge, we created a GPCR screening platform for ligand and drug discovery. This innovative technology enables the cost-effective profiling of ligands and drug compounds against mixtures of hundreds of GPCR-barcoded cell strains in a single experiment. Because a ligand or drug is tested against a collection of receptors all at once, our novel method accelerates the process of identifying potential GPCR ligands and drugs.

## Introduction

G protein-coupled receptors (GPCRs) mediate cellular decision-making and physiological processes by detecting a wide variety of chemical signals, such as small molecule regulators, peptides, and proteins. GPCRs transduce these extracellular signals across the plasma membrane to activate intracellular G proteins that amplify the receptor response through a variety of downstream second messengers (cAMP, IP3, DAG, and Ca^2+^). While more than 360 endoGPCRs comprise the largest and most therapeutically targeted class of membrane receptors in humans, only 30–40% have well-defined biological ligands and are currently druggable (1). The remaining 60–70%, over a hundred of which are classified as “pharmacologically dark” (2), represent an enormous potential for developing new therapeutics and advancing our understanding of GPCR biology.

The yeast *S. cerevisiae* is a useful model for studying human GPCRs (3, 4). Haploid yeast have only one insulated GPCR pathway, known as the pheromone pathway (Figure 1A) (5, 6), that can be adapted to human GPCRs (3, 4). In this system, a human GPCR is coupled to the yeast Gα subunit using C-terminal Gα chimeras in which the last five residues of the yeast Gα is replaced with the last five residues of a human Gα (Figure 1A and S1). Activation of a chimeric GPCR-Gα pair will stimulate a downstream mitogen-activated protein kinase cascade that drives the expression of pheromone-responsive genes. Replacing a dispensable pheromone-responsive gene with a reporter, such as a fluorescent protein, provides a GPCR signaling readout compatible with high-throughput screening formats.

**Fig. 1.**
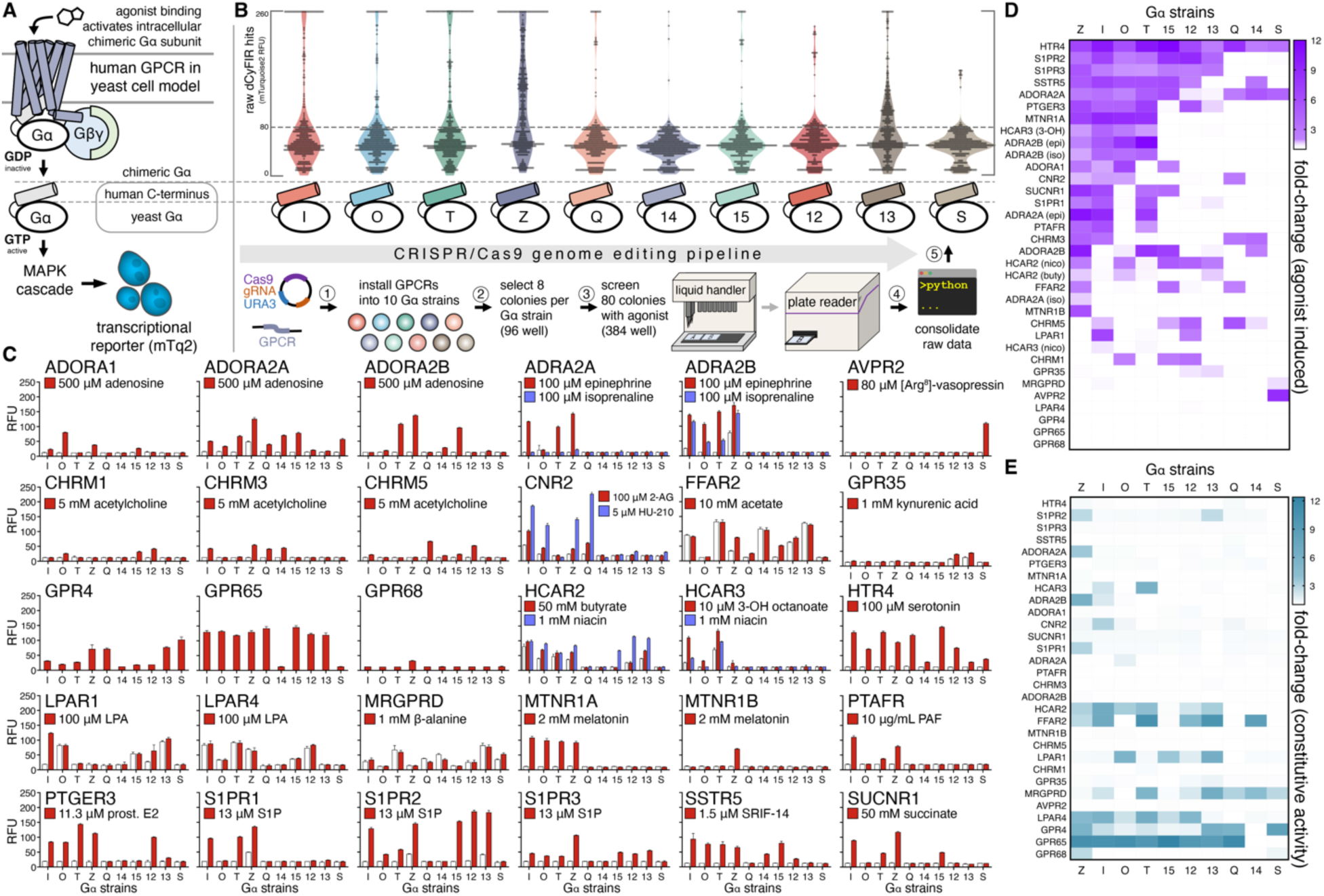
High-throughput CRISPR engineering and DCyFIRscreen validation of the DCyFIR yeast strain library. (A) Simplified schematic of our yeast pheromone pathway model for studying human GPCRs (see Figure S1 for further details). (B) Workflow of our high-throughput CRISPR/Cas9 genome editing pipeline showing primary screening data (more than 5,000 mTq2 fluorescence measurements) for our exploratory panel of 30 human GPCRs. All fluorescence values are reported as relative fluorescence units (RFUs). (C) DCyFIRscreen profiles for 300 GPCR-Gα strains against their known agonists, with error bars representing the SD of n=4 technical replicates. Untreated/treated conditions are represented by white/colored bars. All RFU measurements were quantified using the same gain setting. (D) Heat map of agonist-induced signaling in the 300 GPCR-Gα strains. (E) Heat map of constitutive activity in the 300 GPCR-Gα strains.

Yeast is also an excellent system for high-throughput CRISPR engineering, as genome editing with accuracy, speed, and scale is far easier than in mammalian cells (7–9). This difference in editing efficiency stems from the contrasting mechanisms by which yeast and mammalian cells repair CRISPR/Cas9 targeted double-stranded (DS) DNA breaks. Unlike mammalian cells, which repair DS breaks primarily by non-homologous end joining, yeast repair DS breaks by homologous recombination, readily integrating DNA payloads having sequence homology to CRISPR/Cas9 cut-sites. As such, it is possible to build yeast-based reporter systems that contain genomically stable assay components, preserve available yeast auxotrophies, and obviate heterologous protein over-expression from episomal plasmids.

Here, we report our efforts to build, validate, and apply a multiplexed GPCR profiling platform that exploits the experimental advantages of the yeast pheromone pathway and power of yeast CRISPR. We demonstrate that this discovery platform, which we refer to as Dynamic Cyan induction by Functional Integrated Receptors (DCyFIR), is highly scalable and capable of simultaneously profiling ligands against hundreds of GPCR-Gα combinations with single-cell and single-receptor resolution.

## Results

### High-throughput CRISPR engineering of the DCyFIR yeast strain library

To create the DCyFIR yeast strain library, we first built a panel of 10 Gα reporter strains that covered all possible GPCR-Gα subunit coupling combinations. Each reporter strain contained a pheromone-responsive fluorescent transcription reporter mTq2 (10), a CRISPR-addressable expression cassette in a safe harbor locus on chromosome X, known as X-2 (9), and a unique C-terminal Gα chimera. Additionally, the endogenous GPCR Ste2, GTPase-activating protein Sst2, and cell cycle arrest factor Far1 were deleted from each strain. These deletions were necessary to avoid potential interference from the native yeast GPCR (*ste2*Δ), sensitize the pheromone pathway (*sst2*Δ), and prevent cell cycle arrest upon pathway activation (*far1*Δ). The collection and characterization of the CRISPR deletions, knock-ins, and edits used to build the 10 Gα reporter strains are summarized in Figure S1.

As shown in Figure 1B, our high-throughput CRISPR/Cas9 genome-editing pipeline enabled us to install a set of 30 human GPCRs into all 10 Gα reporter strains, generating all 300 possible GPCR-Gα coupling combinations. We installed each GPCR into the CRISPR-addressable X-2 expression cassette. As a result, each GPCR-Gα strain was barcoded with a genome-integrated GPCR sequence. Although we typically achieved genome-integration efficiencies >80%, we accounted for occasional CRISPR failures by screening 80 candidate colonies for each receptor (8 colonies for each GPCR-Gα combination) against one or more known agonists, or no agonist as in the case of constitutively active receptors such as GPR4, GPR65, and GPR68. As shown in raw CRISPR screening data in Figure 1B, the collection of GPCR-Gα strains exhibited a continuum of strong (> 200 mTq2 relative fluorescence units, RFUs), moderate (> 80 mTq2 RFUs), and weak (30–80 mTq2 RFUs) signaling.

### DCyFIRscreen validation of the DCyFIR yeast strain library

Remarkably, we recapitulated known GPCR agonism in the library of 300 GPCR-Gα yeast strains with 100% accuracy. Our validation process involved two steps. First, we tested each of the 30 sets of 80 candidate GPCR-Gα strains to PCR-verify GPCR genome integration. A single PCR-confirmed hit was selected for each GPCR-Gα reporter strain to establish 10 Gα reporter strains per GPCR. For constitutively active and agonist-inducible GPCR-Gα strains, we selected the best performing (i.e. brightest) PCR-verified colonies. In the second validation step, we used a parallel DCyFIR application, called DCyFIRscreen, to analyze the 300 GPCR-Gα strains in 384-well plate format for known agonism and constitutive activity (Figure 1C). This procedure showed that most GPCRs coupled to multiple Gα chimeras (Figure 1C-E), 24 of 30 GPCRs (80%) coupled to the Gα_z_ chimera (Figure 1D and 1E), and 15 of 30 GPCRs (50%) exhibited some degree of constitutive activity (Figure 1E).

### Developing DCyFIRplex for GPCR-ligand discovery

A major innovation of our DCyFIR strain library is that each strain is barcoded with genome-integrated sequences of a single human GPCR and Gα chimera. This barcoding feature enabled us to develop a second DCyFIR application, called DCyFIRplex (Figure 2A), that we used to profile ligand binding specificity against hundreds of GPCR-Gα strains in a single tube. In the DCyFIRplex method, we monitor agonist-induced mTq2 fluorescence in consolidated mixtures of GPCR-Gα strains using fluorescence activated cell sorting (FACS). Using FACS, we can collect pools of active GPCR-Gα strains (Figures 2A-B) and identify the GPCR barcode(s) that are present using automated quantitative PCR and NanoString deconvolution. As shown in Figure 2A, in a DCyFIRplex experiment equal parts of growth-normalized GPCR-Gα strains are combined with a mRuby3 tracer strain (11) necessary for normalizing different DCyFIRplex runs (Figure 2C) and to empirically determine the optimal duration for FACS (Figure 2D).

**Fig. 2.**
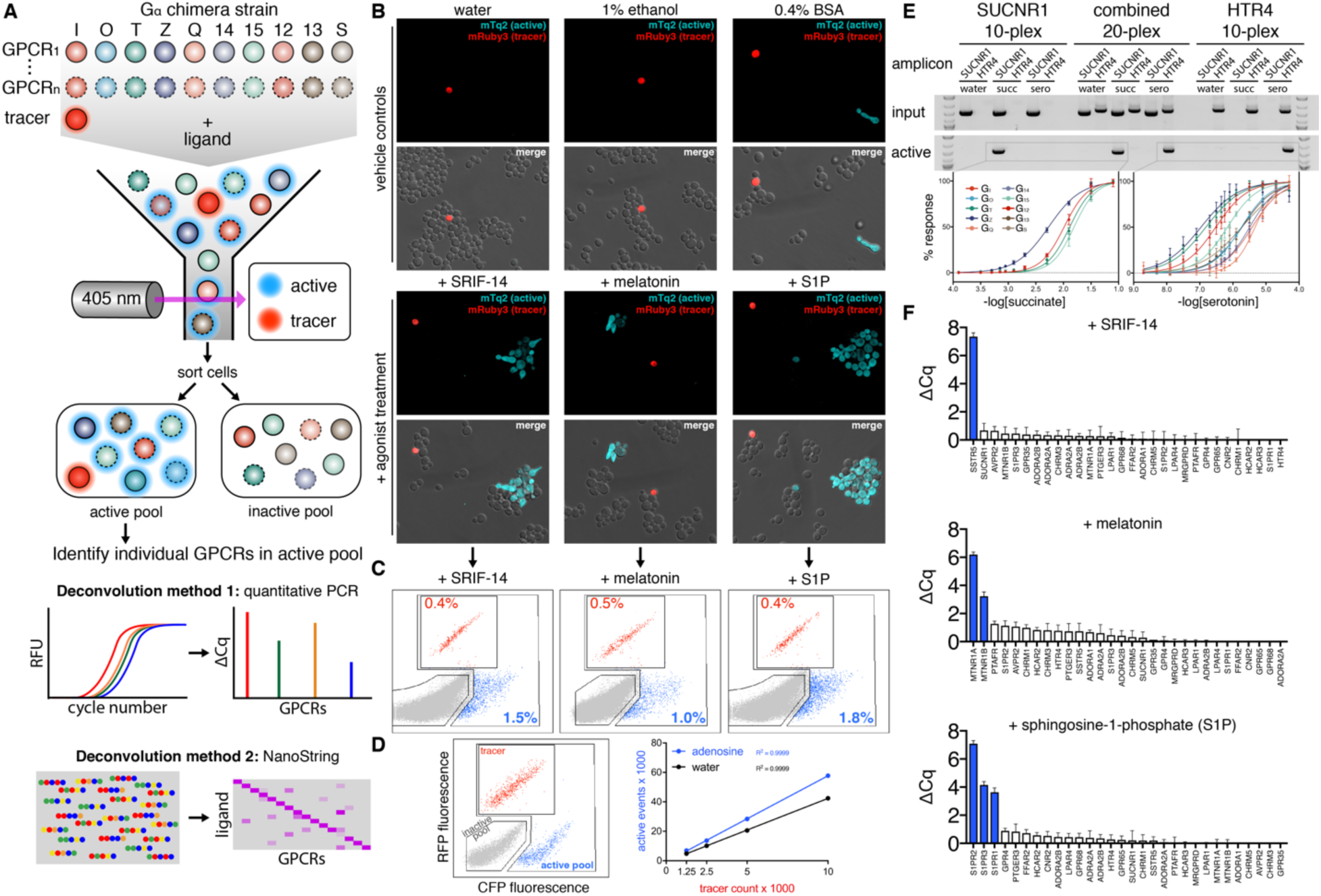
Developing and validating DCyFIRplex. (A) Schematic of the DCyFIRplex workflow showing strain consolidation to build the multiplex, FACS to collect active receptor strain pools, and our two primary multiplex deconvolution techniques. (B) Confocal microscopy images of treated and untreated (vehicle) samples of the GPCR-Gα 300-plex with the added mRuby3 tracer strain (maximum intensity projections, 63X magnification). (C) FACS analysis of the inactive (gray), active (cyan), and tracer (red) pools for the 300-plex shown in panel B. (D) FACS analysis of negative (gray), positive (cyan), and tracer (red) controls (see methods for details); also, a representative standard curve of tracer event counts versus active pool event counts for our reference conditions of -/+ adenosine used to calibrate the FACS sorting procedure. Tracer event counts between 3–5k gave the most consistent deconvolution results. (E) PCR deconvolution of SUCNR1 and HTR4 10-plexes and combined 20-plex visualized by gel electrophoresis; also, the family of normalized titration curves that corresponded to the active GPCR-Gα strains in each 10- and 20-plex (errors bars represent the SD of n=4 technical replicates). (F) DCyFIRplex profiles deconvoluted via qPCR for the agonist-treated 300-plexes characterized in panels B and C. Expected hits are colored blue. ΔCq values correspond to the Cq difference between treated and untreated conditions, with error bars representing the SEM of n=6 repeats derived from 3 independent 300-plex consolidations deconvoluted in technical duplicate. ΔCq values correspond to a log_2_ scale.

The probabilistic character of DCyFIRplex is illustrated in Figures 2B-C. Adding the mRuby3 tracer strain to a 300-plex of GPCR-Gα strains results in a 1 in 301 chance of a tracer sorting event. However, the chance of a sorting event for a given GPCR depends on its number of active GPCR-Gα strains in the multiplex. For example, the melatonin receptor MTNR1A, which signaled in 4 GPCR-Gα strains (Figure 1C), has an expected sorting probability of 4 in 301. As expected, confocal images of untreated samples captured solitary tracer cells surrounded by reporter cells comprising the GPCR-Gα strain pool (Figure 2B). However, we occasionally observed constitutively active GPCR-Gα cells (see Figure 2B, lower right corner of BSA vehicle control). Confocal images taken after agonist treatment showed solitary tracer cells now surrounded by an increased number of active, cyan fluorescent, GPCR-Gα strains (Figure 2B). These active strains tended to form filamentous arrays, a natural process triggered by activation of the pheromone pathway (12). In liquid culture, these cell clusters did not interfere with FACS.

As shown in Figures 2C and 2D, we designed our FACS gates to discern tracer, active, and inactive cell pools. To test this gating strategy, we studied the response of the 300-plex to representative peptide (SRIF-14), small-molecule (melatonin), and lipid (sphingosine-1-phosphate, S1P) agonists that activate one (SSTR5), two (MTNR1A and MTNR1B), and three (S1PR1, S1PR2, S1PR3) receptors, respectively (Figure 2C). Furthermore, we extensively tested the robustness of our gating strategy with panels of negative (the 10 base Gα reporter strains lacking GPCRs) and positive controls (two strains that constitutively expressed the mRuby3 tracer or mTq2 reporter) (Figure 2D). We typically used 3–5k tracer counts to collect an active pool of 20–25k sorting events (Figure 2D). As shown in Figure 2C, only a small fraction of sorting events corresponded to tracer (0.4, 0.5, and 0.4%) and active (1.5, 1.0, 1.8%) cell pools following agonist treatments. Having demonstrated our ability to sort activated GPCR-Gα strains, we were poised to advance the speed, scope, and scale of GPCR profiling using DCyFIRplex.

### DCyFIRplex profiling identifies new interactions for known agonists

To demonstrate the DCyFIRplex method, we first identified GPCR gene(s) sorted into active pools using a 20-plex of SUCNR1 and HTR4 GPCR-Gα strains (Figure 2E). As expected, treatment of the 20-plex with succinate and serotonin resulted in active pools that exclusively contained barcoded gene sequences for the receptors SUCNR1 or HTR4, respectively. Following on this success, we expanded our experiments to include all 300 GPCR-Gα strains. Representative results for the 300-plex treated with agonists SRIF-14 (SSTR5), melatonin (MTNR1A and MTNR1B), and S1P (S1PR1, S1PR2, S1PR3) are shown in Figure 2F. As anticipated, each agonist treatment resulted in active pools exclusively containing the expected receptor gene(s) (Figure 2F). To demonstrate robustness, we successfully repeated the DCyFIRplex procedure for our full set of agonists (Figures 3A-B), again recapitulating known GPCR agonism with 100% accuracy.

**Fig 3.**
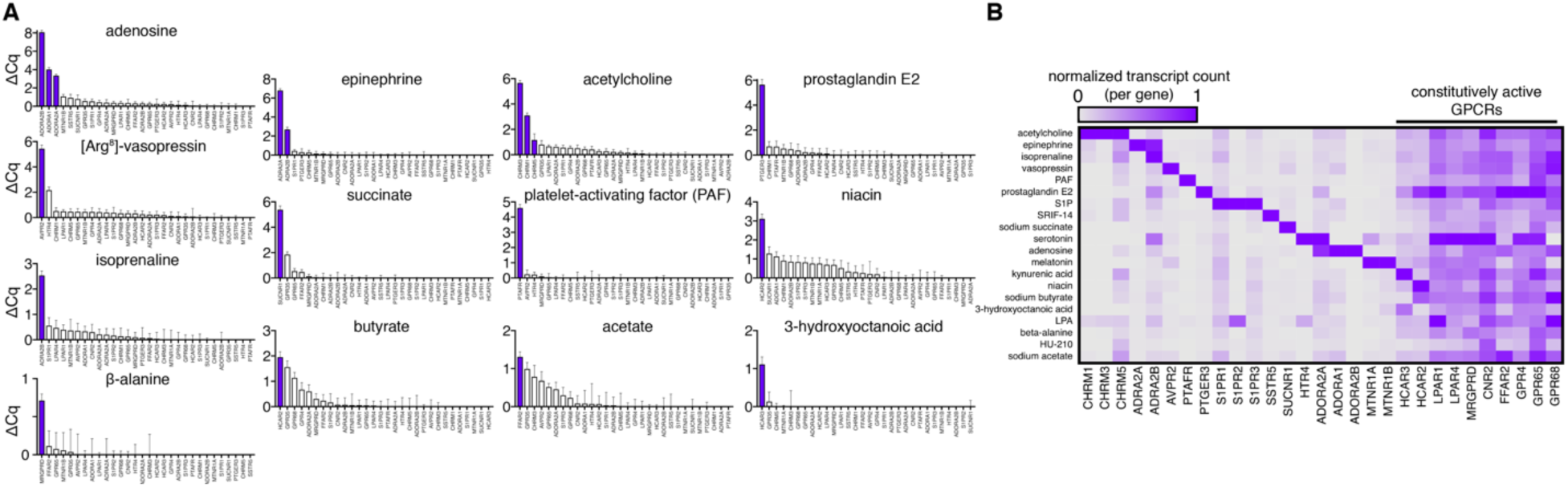
Using DCyFIRplex profiling to recapitulate known agonist interactions. (A) DCyFIRplex profiles for known GPCR agonists in our 30-receptor panel deconvoluted via qPCR. (B) Same samples as in panel A deconvoluted using NanoString. (A, B) ΔCq values correspond to the Cq difference between treated and untreated conditions, with error bars representing the SEM of n=6 repeats derived from 3 independent 300-plex consolidations deconvoluted in technical duplicate. ΔCq values correspond to a log_2_ scale. NanoString transcript counts were collected in technical duplicate, averaged, and normalized to the maximal RNA transcript count for each GPCR gene.

The DCyFIRplex experimental format also led us to discover new interactions for known agonists (Figure 4A), including lysophosphatidic acid (LPA) with S1PR2, serotonin with MTNR1A, and the metabolite kynurenic acid (KYNA) with both ADRA2B and the dark receptor HCAR3. As shown in Figure 4B, we used DCyFIRscreen to confirm each new interaction. In only one case (KYNA), did we observe a reduction in signaling upon ligand treatment (ADRA2B, Figure 4A), suggesting that KYNA was an inverse agonist or negative allosteric modulator of ADRA2B. To investigate these possibilities and validate our discoveries, we performed detailed titrations that quantified micromolar interactions of serotonin with MTNR1A and ADRA2B (Figure 4C), KYNA with HCAR3 and ADRA2B (Figure 4D), and LPA with S1PR2 (Figure S2). Additional HCAR3 and ADRA2B titrations with their known ligands (3-hydroxyoctanoic acid and epinephrine) in the presence of KYNA demonstrated that KYNA is an orthosteric agonist of HCAR3 (Figure S2) and a negative allosteric modulator of ADRA2B (Figure 4D). Lastly, head-to-head KYNA titrations of GPR35 and HCAR3 revealed that KYNA has a nearly 20-fold higher affinity for HCAR3 (EC_50_ 41 μM) compared to its known target, GPR35 (EC_50_ 795 μM) (Figure 4D).

**Fig. 4.**
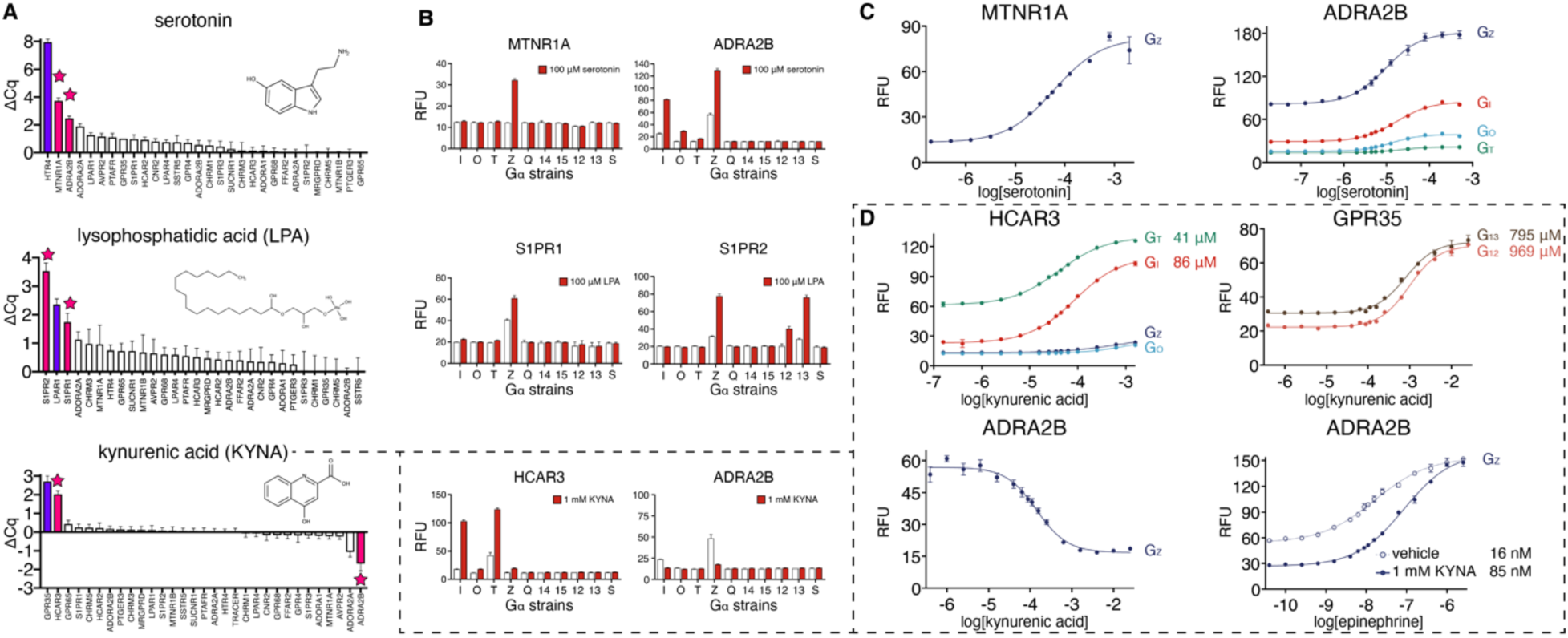
Using DCyFIRplex to discover new interactions for known GPCR agonists. (A) DCyFIRplex profiles identifying new GPCR-ligand interactions (pink bars) discovered in the process of screening known agonists (purple bars) within our panel of 30 exploratory receptors. (B) DCyFIRscreen profiles confirming the DCyFIRplex discoveries in panel A, with error bars representing the SD of n=4 technical replicates. (C, D) Select titrations confirming the DCyFIRplex discoveries in panel A, with error bars representing the SD of n=4 technical replicates. Dashed lines and boxes indicate datasets showing that KYNA activates HCAR3 with greater potency than GPR35 and is also an endogenous negative allosteric modulator of ADRA2B. Full titration datasets are available in Figure S2.

### DCyFIR profiling of 320 metabolites finds new GPCR agonists and allosteric modulators

To demonstrate scalability of the DCyFIR platform, we next profiled the 300-plex of GPCR-Gα strains against a library of 320 endogenous human metabolites using a 6-step procedure (Figure 5A). As shown in Figure 5B, we divided the 300-plex into three sets. The GPCR-Gα strains comprising sets 2 (100-plex) and 3 (90-plex) exhibited lower constitutive activity than those in set 1 (110-plex). We classified potential metabolite hits as having a Z-score > 1 (Figure 5B) and fluorescence microscopy images with marked increases in mTq2 fluorescence, filamentous arrays, and shmooing (Figure 5C). Using these criteria, we recovered known metabolite agonists for GPCRs in our exploratory panel, such as adenosine, melatonin, prostaglandin E2 (Figure 5B). However, most metabolite hits corresponded to new interactions, including tryptamine and dopamine agonism of adrenergic receptors (Figure 5D), phenylethanolamine (PEOA) agonism of ADRA2B (Figure 6A), and petroselinic acid agonism of S1PR1, S1PR2, and LPAR1 (Figure 6A).

**Fig. 5.**
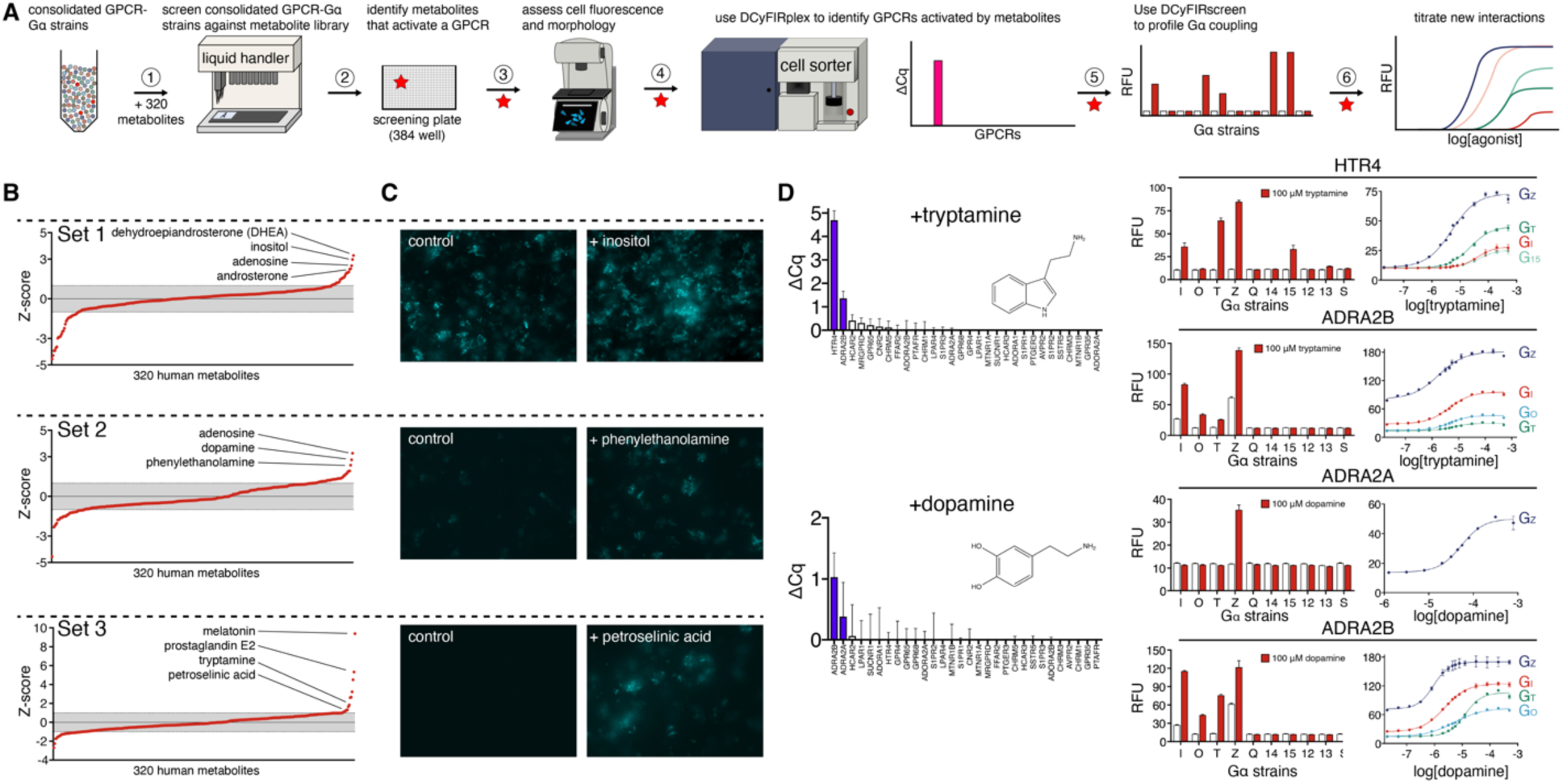
DCyFIR profiling of a human metabolite library. (A) Step-by-step workflow used to screen a library of 320 endogenous human metabolites. (B) Z-score profiles for metabolite screens of receptor set 1 (ADORA1, ADORA2A, FFAR2, GPR4, GPR65, GPR68, HCAR2, HCAR3, LPAR1, LPAR4, MRGPRD), set 2 (ADORA2B, ADRA2A, ADRA2B, AVPR2, CHRM1, CHRM3, CHRM5, CNR2, GPR35), and set 3 (HTR4, MTNR1A, MTNR1B, PTAFR, PTGER3, S1PR1, S1PR2, S1PR3, SSTR5, SUCNR1). Grey bands indicate Z-scores between ± 1. (C) Representative fluorescence microscopy images for Z-score hits in receptor subsets 1 (110-plex), 2 (90-plex), and 3 (100-plex). (D) Discovery workflow illustrating tryptamine agonism of HTR4 and ADRA2B and dopamine agonism of ADRA2A and ADRA2B. Once tryptamine and dopamine were identified as hits (steps 1-3 in panel A), we used DCyFIRplex profiling to identify their GPCR target(s), DCyFIRscreen profiling to identify their Gα coupling pattern(s), and titrations to quantify their EC_50_ values (steps 4-6 in panel A).

**Fig. 6.**
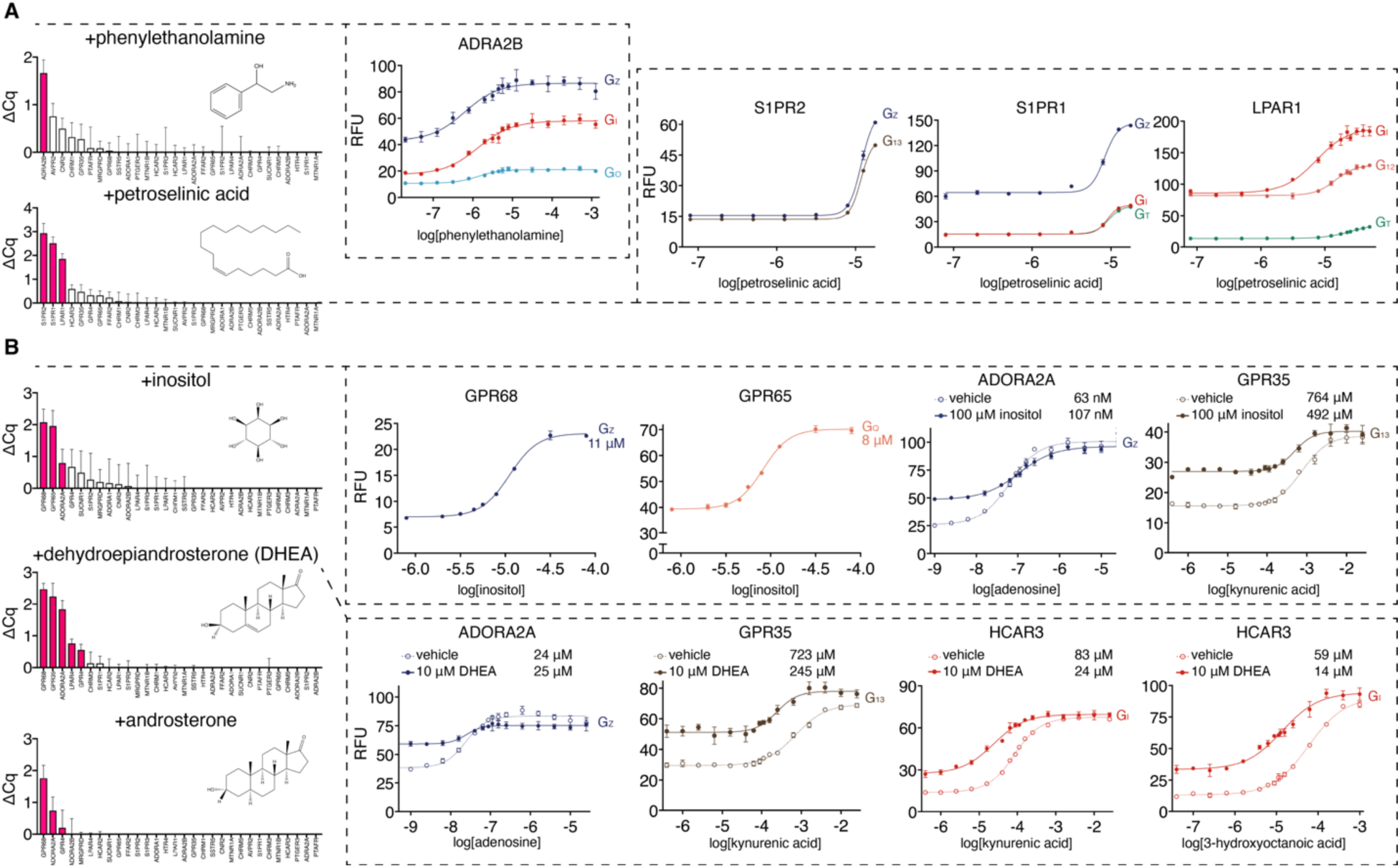
Identification and validation of new GPCR-metabolite interactions. (A) DCyFIRplex profiles and titrations for new metabolite agonists and (B) positive allosteric modulators. For panel B, full titration datasets for all GPCR-Gα coupling combinations are available in Figure S4. DCyFIRplex error bars represent the SEM of n=6 repeats derived from 3 independent 300-plex consolidations deconvoluted in technical duplicate, and titration error bars represent the SD of n=4 technical replicates.

DCyFIR profiling also enabled us to discover metabolites that function as allosteric modulators. Remarkably, most of these findings involved dark receptors GPR4, GPR65, GPR68, and HCAR3. As shown in Figure 6B, inositol and the steroid metabolites DHEA and androsterone interacted with multiple prototypic and understudied GPCRs in the DCyFIRplex experiment. To rigorously assess the specificity of these ligands, we DCyFIRscreen profiled inositol, DHEA, and androsterone against many GPCRs (>150 control experiments are available in Figures S3-S6). This vetting process indicated that inositol modulated 5 receptors (ADORA2A, CNR2, GPR35, GPR65, GPR68), and that the structurally similar steroid metabolites DHEA and androsterone modulated 8 receptors (ADORA2A, GPR35, GPR4, GPR68, HCAR2, HCAR3, LPAR1, LPAR4). Interestingly, all three metabolites modulated a trio of GPCRs (ADORA2A, GPR35, and GPR68). Based on these observations, we speculated that inositol, DHEA, and androsterone were broad-spectrum allosteric modulators. As show in Figure 6B, we tested this idea by titrating select receptors with inositol (GPR65, GPR68) or orthosteric agonists in the presence of inositol and DHEA (ADORA2A, GPR35, HCAR3). These experiments demonstrated that inositol and DHEA are negative allosteric modulators of ADORA2A, inositol is a positive allosteric modulator of GPR65, GPR68, and GPR35, and DHEA is a positive allosteric modulator of HCAR3 and GPR35.

## Discussion

The objective of this study was to demonstrate the feasibility and potential of our innovative DCyFIR platform. Using this new technology, we have identified a relatively large number of novel GPCR-ligand interactions by profiling a relatively small number of compounds and receptors. These discoveries are solely attributable to the multiplex format of the DCyFIR experiment, which enables simultaneous ligand profiling of many GPCRs at once. In the discussion that follows, we characterize a select number of our findings in the context of their potential biological relevance.

### New interactions between GPCRs and amino acid metabolites

In this work we showed that the DCyFIR platform can be used to identify new interactions between GPCRs and small molecule regulators. For example, we identified several new GPCR-ligand interactions involving three amino acid metabolites that belong to the tryptophan pathway: KYNA, serotonin, and tryptamine. Of these findings, our discovery that KYNA is a more potent agonist of HCAR3 than GPR35 is particularly notable. KYNA is known to play important roles in neuroprotection, depression, schizophrenia, obesity, diabetes, and cancer (13–15). Prior to this study, the only GPCR known to respond to KYNA was GPR35 (16). However, we have shown that KYNA activates HCAR3 with a nearly 20-fold lower EC_50_ than GPR35. Interestingly, we also found that similarities between HCAR3 and GPR35 extended beyond KYNA agonism, as both receptors exhibited PAM interactions with the steroid metabolite DHEA. In the case of HCAR3, DHEA PAM interactions extended to both new (KYNA) and known (3-hydroxyoctanoic acid) orthosteric agonism, similar to the dual regulation imposed by KYNA and DHEA on the N-methyl-D-aspartate (NMDA) receptor (17). Based on these observations, we believe a comprehensive re-assessment of HCAR3, GPR35, and KYNA agonism in relevant cell types and physiological systems is warranted.

### Cross-activation of lipid-binding GPCRs

Our set of 30 GPCRs included members of two evolutionarily-related lipid receptor families, LPAR and S1PR. Both receptor families are involved in inflammatory responses, fibrosis, and a variety of other disorders (18, 19). They also share similar spatially-conserved residues that determine their respective specificities for LPA and S1P metabolites (20). Here we confirmed the known cross-activation of LPA for S1PR1 (21, 22), and to the best of our knowledge report the first LPA cross-activation of S1PR2. Consistent with previous LPA studies, we found that LPA activated both S1PR1 and S1PR2 with higher EC_50_ values than LPAR1, indicating that S1PR1 and S1PR2 are low affinity LPA receptors. Although it has been suggested that the micromolar LPA concentrations needed to activate S1PR1 and S1PR2 receptors may not be biologically relevant (21), local (paracrine and autocrine) and systemic (endocrine) metabolite concentrations can regularly occur at micromolar levels in response to biological cues. For example, the succinate receptor SUCNR1 has a low affinity for succinate, however, the accumulation of succinate under stress conditions, such as ischemia, activates SUCNR1 (23, 24). Interestingly, we did not observe LPA cross-activation of S1PR3.

### Allosteric modulators of understudied GPCRs

The sensitivity of the DCyFIR platform was demonstrated by our discovery of several allosteric regulators for prototypic and understudied GPCRs. Although most of these interactions had micromolar affinity, they were repeatedly detectable in both DCyFIRscreen and DCyFIRplex experimental modes. Remarkably, this sensitivity even extended to GPCR-ligand interactions having near-millimolar affinity, such as the EC_50_ of KYNA for GPR35. Given that allosteric modulators are often attractive drug candidates, we believe that this novel feature of DCyFIR profiling will enable the identification of chemical leads over a wide range of affinities, greatly expediting the process of developing new pharmacological tools and drugs.

Notably, all of the allosteric modulators we discovered in our metabolite screen interacted with pharmacologically dark GPCRs. Inositol, a structural isoform of glucose, was a PAM of GPR65 and GPR68, while DHEA, the most abundant circulating steroid hormone in humans (25), was a PAM of GPR4, GPR68, and HCAR3. To the best of our knowledge, this is the first report of endogenous PAMs for these understudied receptors. Furthermore, there has been a long-standing interest in discovering mediators of steroid responses outside the nucleus (26). Our finding that DHEA and androsterone act as PAMs of several prototypic and dark GPCRs, suggests that there are areas of steroid metabolism waiting to be explored. Lastly, our steroid-related discoveries showcase one of the major advantages of our yeast-based platform: because yeast lack cholesterol and have primitive steroid pathways (27), there is little to no steroid interference from the model system or its genetic background in the discovery process. Based on our DHEA and androsterone findings, we are currently exploiting this feature of the yeast system to comprehensively examine the unexplored frontier of GPCR-steroid pharmacology.

### The potential of the DCyFIR platform

In conclusion, we have demonstrated the utility of the DCyFIR platform for simultaneously profiling many GPCRs against individual ligands and chemical libraries. Furthermore, we established that this innovative approach facilitates the rapid, inexpensive, and comprehensive assessment of ligand specificity and receptor promiscuity. We believe that combining the DCyFIR platform with other GPCR screening approaches has the potential to open new avenues in GPCR research. By quickly and openly sharing the results of such studies, the longer-term process of validating the biological and clinical relevance of new GPCR-ligand interactions, as well as developing lead compounds into future drugs, can be expedited within the community. As our library of GPCR-Gα strains continues to grow, so too will the scale of our DCyFIR profiling capabilities and ligand discovery efforts.

## Methods

### Media

The different media types used in this study are detailed in the Table S1.

### Plasmids

All plasmid vector sequences are provided in Supplementary Dataset 3.

### Strains

The 323 yeast strains created and used in this work are listed in Supplementary Dataset 1.

### Engineering CRISPR-optimized yeast strains for human GPCR studies

The collection of 323 yeast strains created and used in this study are detailed in Supplementary Dataset 1. The major steps we used to build the 10 base GPCR-Gα strains are summarized below. We used CRISPR to make every gene deletion, replacement, edit, and knock-in in this study (see “*CRISPR transformation reaction”* method for further detail). See Supplementary Dataset 3 for a complete list of CRISPR plasmids.

#### Deletion of signaling components to sensitize the pheromone pathway

In the first steps of our strain engineering efforts, we created our DI2Δ strain by sequentially deleting the pheromone pathway components FAR1 and SST2. The factor arrest protein (FAR1) was deleted to prevent cell-cycle arrest upon pathway activation and the GTPase-activating protein (SST2) was deleted to sensitize the pheromone pathway by prolonging Gα activation. Our CRISPR gene deletion procedure employed two CRISPR vectors, pML107 and pT040, each having their own selectable markers LEU and URA. Vector pT040 contained a guide RNA sequence that targeted the N-terminal/C-terminal region of the gene to be deleted. These vectors were co-transformed with DNA payload comprising homology arms generally having 60-100 bp of sequence immediately upstream and downstream of the targeted open reading frame.

#### Installation of the mTq2 transcriptional reporter

Following the creation of the DI2Δ strain, we replaced the pheromone-responsive gene *FIG1* open reading frame with the cyan fluorescence protein mTq2. As with our CRISPR gene deletion procedure, we replaced the FIG1 open reading frame with the mTq2 gene using two CRISPR vectors pML107 and pT040, each having their own selectable markers LEU and URA. Vectors pML107 and pT040 contained a guide RNA sequence that targeted the N-terminal/C-terminal region of the *FIG1* gene. These vectors were co-transformed with DNA payload comprising homology arms having 60 bp of sequence immediately upstream and downstream of FIG1 open reading frame. The resultant genotype of this strain, which we refer to as DI2Δ fig1Δ::mTq2, was BY4741 far1Δ sst2Δ fig1Δ::mTq2.

#### Deleting the endogenous yeast GPCR STE2 gene

Following the creation of the DI2Δ fig1Δ::mTq2 strain we deleted the native yeast GPCR gene (STE2) using the same plasmids and procedure described in the section *Deletion of signaling components to sensitize the pheromone pathway.* This new strain, DI3Δ fig1Δ::mTq2, had the genotype BY4741 far1Δ sst2Δ ste2Δ fig1Δ::mTq2.

#### Installation of the X-2 landing pad

Following the creation of the DI3Δ fig1Δ::mTq2 strain, we installed our CRISPR-addressable expression cassette (see “*Creating the CRISPR-addressable expression cassette into the X-2 locus*” for details) into the X-2 locus of chromosome X. To install the X-2 landing pad into the X-2 safe harbor locus, we PCR-amplified the landing pad sequence from the pMARQ vector and co-transformed the resultant DNA payload with the CRISPR vector pML104 X2 The resultant genotype of this strain, which we refer to as DI3Δ fig1Δ::mTq2 P1, was BY4741 far1Δ sst2Δ ste2Δ fig1Δ::mTq2 X-2:P_TEF1a_-UnTS-T_CYC1b_.

#### Genome-editing to create humanized yeast C-terminal Gα chimeras

To build our panel of 10 GPCR-Gα base reporter strains, we created 10 different versions of our DI3Δ fig1Δ::mTq2 P1 strain, each having its own unique Gα C-terminal yeast/human chimera. In each Gα chimera, the last five yeast residues of the yeast Gα subunit, Gpa1, were replaced by the last five residues of a human Gα subunit (see Figure S1 and Table S1 for sequence details). Due to C-terminal degeneracies, all 16 human Gα genes could be represented by 10 Gα C-terminal chimeras. A codon-optimized DNA payload for each Gα chimeric sequence was designed as a gBlock gene fragment (Integrated DNA Technologies) comprising the 15 bp sequence of a human Gα C-termini flanked by 123 bp homology arms that targeted the C-terminus of the yeast Gα subunit sequence. These synthetic DNA payloads were co-transformed with the CRISPR vectors pML107 and pT040 GPA1:1373, each having their own selectable markers LEU and URA. The resultant genotypes of these strains, which we refer to as DI DCyFIR P1 I, DI DCyFIR P1 O, DI DCyFIR P1 T, DI DCyFIR P1 Z, DI DCyFIR P1 Q, DI DCyFIR P1 14, DI DCyFIR P1 15, DI DCyFIR P1 12, DI DCyFIR P1 13, and DI DCyFIR P1 S, are described in Supplementary Dataset 1.

#### Installation of human GPCRs into X-2 CRISPR-addressable expression cassette

All human GPCR DNA sequences were sourced from the Presto-TANGO plasmid library (28), using primers to PCR amplify only the GPCR open reading frame, avoiding the additional N- and C-terminal DNA sequence elements in the Presto-TANGO plasmid constructs. Using two rounds of PCR amplification, we extended each GPCR sequence with homology arms corresponding to sequences within the TEF promoter and CYC1b terminator of the X-2 landing pad. For the first round of PCR, we used receptor-specific primers having ∼45 bp homology overhangs. In a second round of PCR, we used universal primers to extend both homology arms to a final length of 60 bp. All primer and GPCR sequences are listed in Supplementary Datasets 2 and 4. With the exception of the muscarinic receptors CHRM1, CHRM3, and CHRM5, native GPCR sequences were used (i.e. no affinity tags or localization sequences were added). However, residues corresponding to the third intracellular loop (iL3) of CHRM1, CHRM3, and CHRM5 were deleted to reproduce iL3 deletion results that were previously published in a similar yeast system (29). As with these original studies, we found that full-length CHRM1, CHRM3, and CHRM5 did not functionally express in our system. The CHRM1, CHRM3, and CHRM5 sequences in Table S1 correspond to the iL3 loop deletion variants. To install human GPCRs into the X-2 landing pad, the amplified GPCR PCR product with 60 bp homology arms was co-transformed with the CRISPR vector pML104 X2 UnTS using the approach described in “*CRISPR transformation reactions*”. Because we installed each human GPCR into all 10 base GPCR-Ga reporter strains, we produced a library of 300 new GPCR-Gα strains barcoded with a human GPCR.

### DCyFIRscreen protocol

Individual Gα reporter strains were grown in SCD LoFo pH = 7.0 at 30 °C to an OD = 1.0 in a 2.0 mL 96-well DeepWell block (Greiner; 780271-FD). Cells were normalized to an OD of 0.1 in SCD LoFo pH = 7.0 using a Biomek NXp liquid-handling robot. 10X ligand/vehicle stocks were prepared (see Key Resource Table) and 4 μL were distributed to each well of a 384-well plate (Greiner; 781096) in quadruplicate using a Biomek NXp. 36 μL normalized cells were distributed to each well containing the appropriate 10X ligand/vehicle. Plates were sealed with a breathable cover (Diversified Biotech; BERM-2000) and incubated at 30 °C. Fluorescence readings were collected after 18 hours using a plate reader (ClarioStar, BMG LabTech, Offenburg, Germany) (bottom read, 10 flashes/well, excitation filter: 430-10 nm, dichroic filter: LP 458 nm, emission filter: 482-16 nm, gain = 1300 (1500 for Figure 1B.)). Absorbance readings were also collected after 18 hours using the same instrument (22 flashes/well, excitation filter: 600 nm).

### DCyFIRplex protocol

Control/tracer strain(s) (DI P1 mTq2, individual Gα reporter strains lacking an integrated receptor, DI P1 mRuby3, and the 300 GPCR-Gα strains were grown in SCD LoFo pH = 5.0 to saturation in individual wells of a 2.0 mL 96-well DeepWell block (Greiner; 780271-FD). The 10 GPCR-Gα strains for a single receptor were then consolidated in growth-normalized amounts into single wells of a DeepWell block using a Biomek NXp (each well is comprised of one unique receptor in all 10 Gα reporter strains). Each receptor-consolidated well and control well was grown to mid-log phase in SCD LoFo pH = 7.0 then further consolidated into a single tube (300-plex; 30 receptors, 300 Gα strains) in growth-normalized amounts (this consolidation was performed three times for each sorting procedure [n = 3]). 10X ligand/vehicle stocks were added to individual wells of a DeepWell block. The consolidated 300-plex or control strains were then added to each well and grown in the presence of ligand/vehicle overnight, so that each culture would reach an OD = 4.0 before DCyFIRplex profiling. Samples were washed with sterile ddH_2_O and normalized to an OD = 2.0 in SCD LoFo pH = 7.0. Tracer cells were added to each sample at a 1:301 ratio. The final mixture was then transferred into a glass sample tube (USA Scientific; 1450-2810) and used for cell sorting. A BD FACSAria-II cell sorter was used for all DCyFIRplex experiments to assess mTq2 (405 nm excitation, 450/50 nm emission) and mRuby3 fluorescence (535 nm excitation, 610/20 nm emission). A gating strategy was set using the three control samples (DI P1 mTq2, individual Gα reporter strains lacking an integrated receptor, and DI P1 mRuby3) such that tracer cells and any cell expressing mTq2 was sorted into a 14 mL collection tube (USA Scientific; 1485-2810) containing 500 μL YPD. Samples treated with water or 500 μM adenosine (well-characterized using DCyFIRscreen, inexpensive, and water-soluble) were used to build a standard curve measuring total events in the mRuby3 and mTq2 positive gates. The standard curve from a water-treated 300-plex was used to determine the number of tracer events that would correspond to 15,000 events in the mTq2 gate for a water-treated 300-plex. Each sample was sorted until the standardized tracer count was reached. Sorted cells were enriched by outgrowth in 5 mL YPD at 30°C with shaking (200 rpm) for 18 hours. Cells were harvested by centrifugation at 3,000xg for 5 min, and resuspended in 1 mL ddH_2_O. Cells were either processed immediately for qPCR deconvolution or frozen in 100 μL aliquots for storage at −20 °C. The set of samples comparatively deconvoluted by qPCR and NanoString methods were derived from aliquots of the same DCyFIRplex experiments (see Supplementary Methods for details).

## Supporting information

Data Set 1

Data Set 2

Data Set 3

Data Set 4

## Acknowledgements

We would like to thank Dr. Antonio Barrientos, Dr. Grace Zhai, and Dr. Vladlen Slepak for their critical review of this manuscript. This work was supported by the NIH through the NIGMS (R35GM119518) and NIH Common Fund for Illuminating the Druggable Genome (R03TR002908) to D.G.I.

## Supplementary Information

**Fig. S1.**
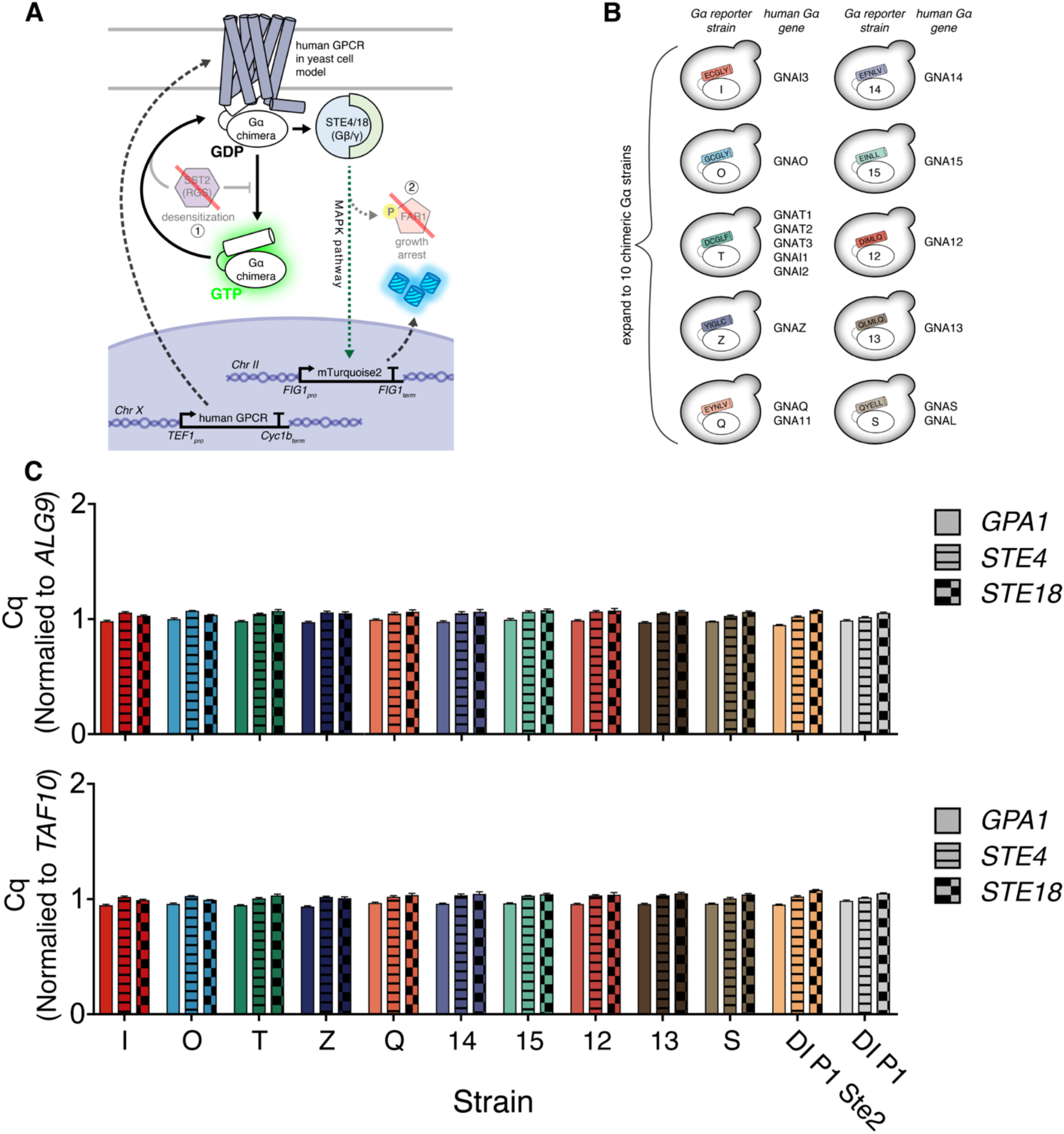
Engineering and characterizing the base GPCR-Gα reporter strains. (A) Schematic of the engineered yeast pheromone pathway for studying human GPCRs. A gene encoding a human GPCR is directly integrated into the yeast genome into a synthetic expression cassette at the X-2 locus. Human GPCR signaling through a chimeric yeast Gα protein is prolonged due to the deletion of a negative regulator (RGS protein, SST2). MAP kinase cascade signaling drives the expression of a bright cyan fluorescent protein, mTq2, that was directly installed into the yeast genome replacing a pheromone-responsive gene, FIG1. Deletion of FAR1 prevents cell cycle arrest upon MAPK signaling. (B) The 16 genes encoding human Gα proteins can be represented by 10 degenerate Gα chimeras (10 unique yeast strains), each with the corresponding 5 C-terminal amino acids from humans. (C) qPCR data illustrating similar heterotrimeric G protein expression levels across all 10 GPCR-Gα reporter strains, Ste2 rescue strain (DI P1 Ste2), and base reference strain (DI P1). Data reported as Cq values normalized to two housekeeping genes (ALG9 and TAF10) with error bars representing SEM of n=3-5 technical replicates.

**Fig. S2.**
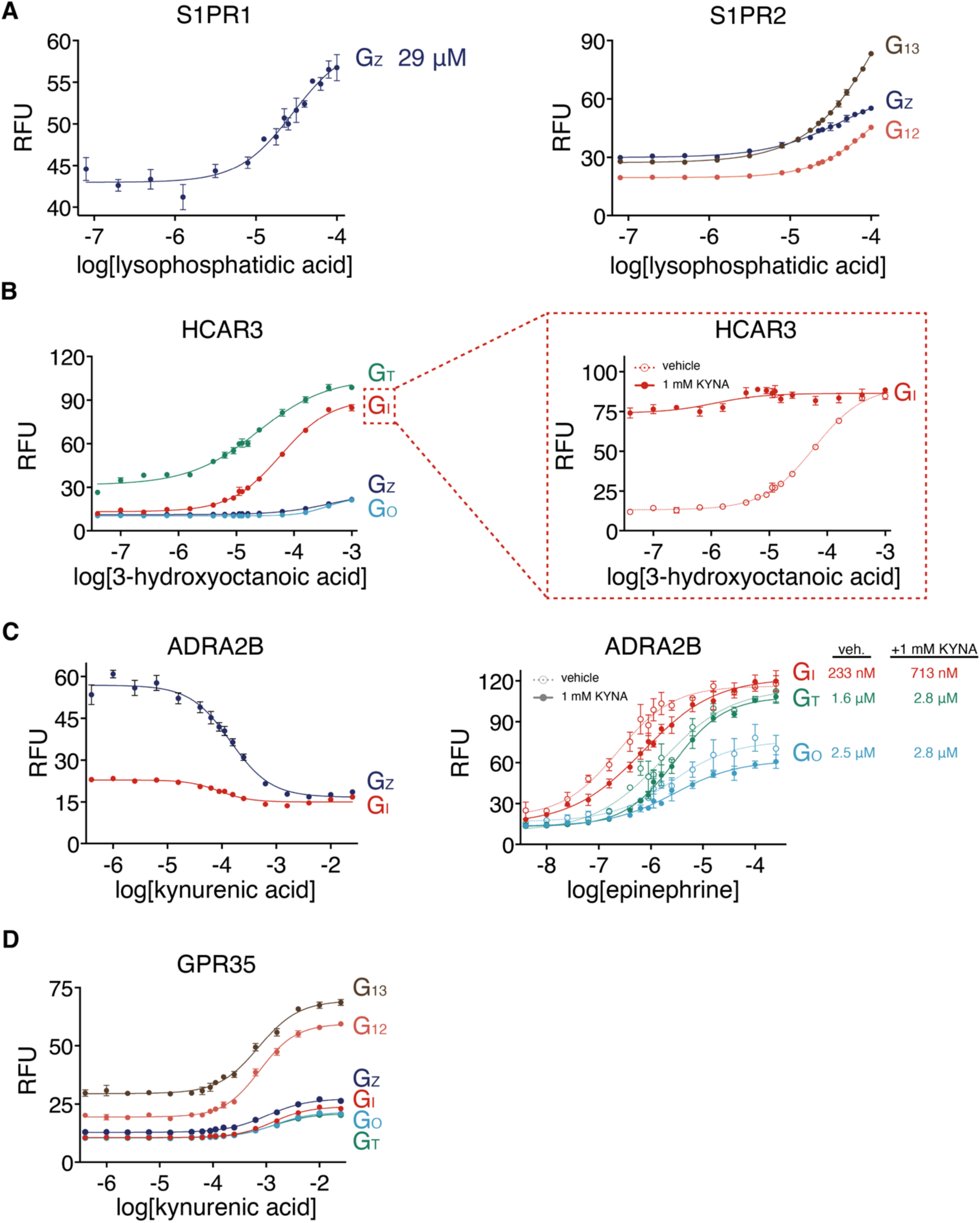
Follow-up titrations and control experiments for new ligand discoveries. (A) Dose-response curves for S1PR1 and S1PR2 with lysophosphatidic acid (LPA). Error bars represent SD of n=4 technical replicates. (B) Dose-response curves for HCAR3 with 3-hydroxyoctanoic acid (left) and 3-hydroxyoctanoic acid (dashed line, open circles) in the presence of kynurenic acid (solid line, closed circles) (right). Error bars represent SD of n=4 technical replicates. (C) Dose-response curves for ADRA2B with kynurenic acid (left) and epinephrine (dashed line, open circles) in the presence of kynurenic acid (solid line, closed circles) (right). Error bars represent SD of n=4 technical replicates. (D) Dose-response curves for GPR35 with kynurenic acid. Error bars represent SD of n=4 technical replicates.

**Figure S3.**
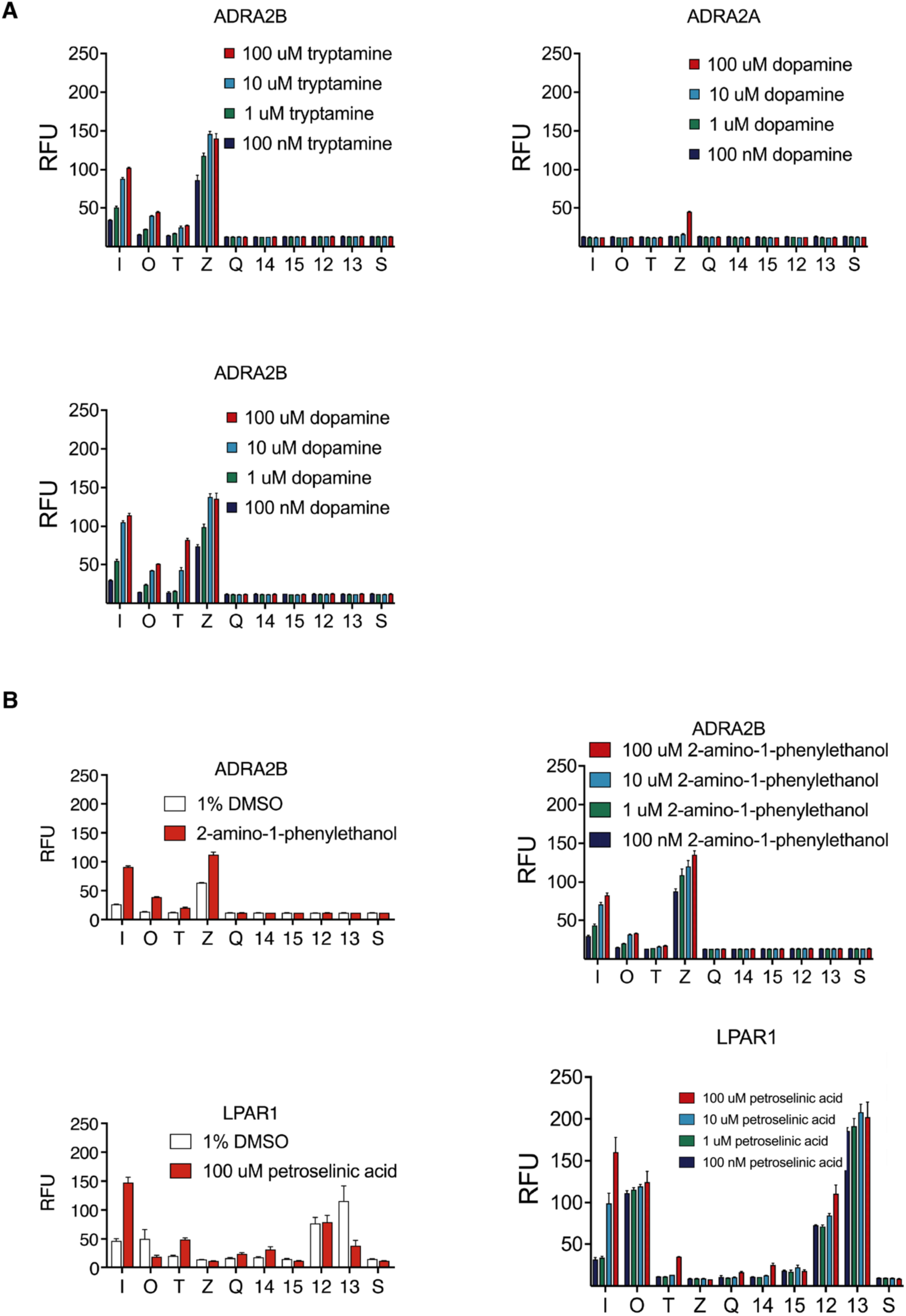
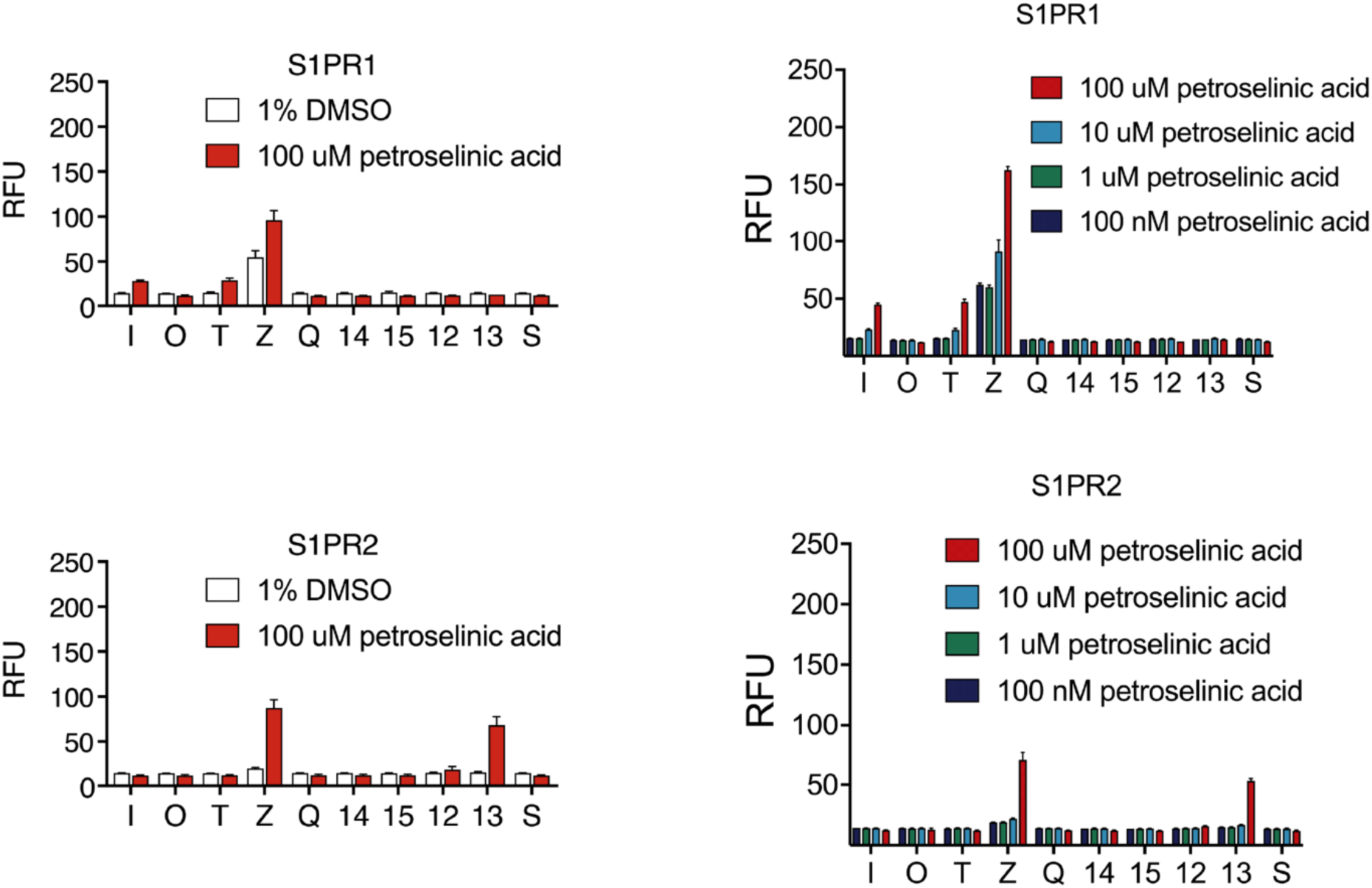
DCyFIRscreen profiles and coarse titrations to confirm new metabolite agonists and allosteric modulators. (A) Coarse 4-point titrations for ADRA2B with tryptamine and dopamine, and ADRA2A with dopamine. Error bars represent SD of n=4 technical replicates. (B) DCyFIRscreen profiles and coarse 4-point titrations for new metabolite interactions (2-amino-1-phenylethanol and petroselinic acid). Error bars represent SD of n=4 technical replicates.

**Fig. S4.**
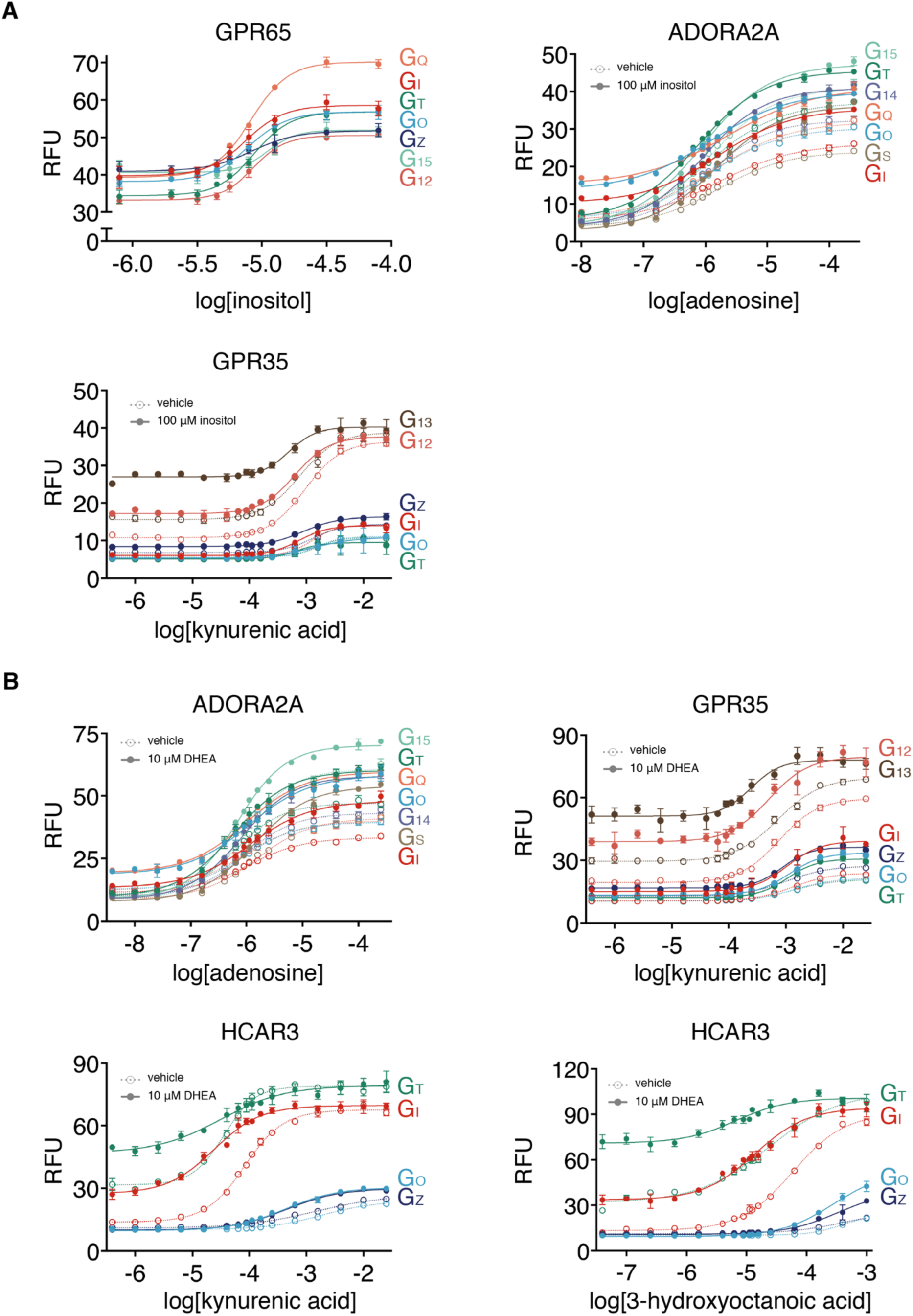
Detailed titrations for new metabolite allosteric modulators. (A) Dose-response curves for GPR65, ADORA2A, and GPR35 (dashed lines, open circles) in the presence of inositol (solid lines, closed circles). Error bars represent SD of n=4 technical replicates. (B) Dose-response curves for ADORA2A, GPR35, and HCAR3 (dashed lines, open circles) in the presence of DHEA (solid lines, closed circles). Error bars represent SD of n=4 technical replicates.

**Fig. S5.**
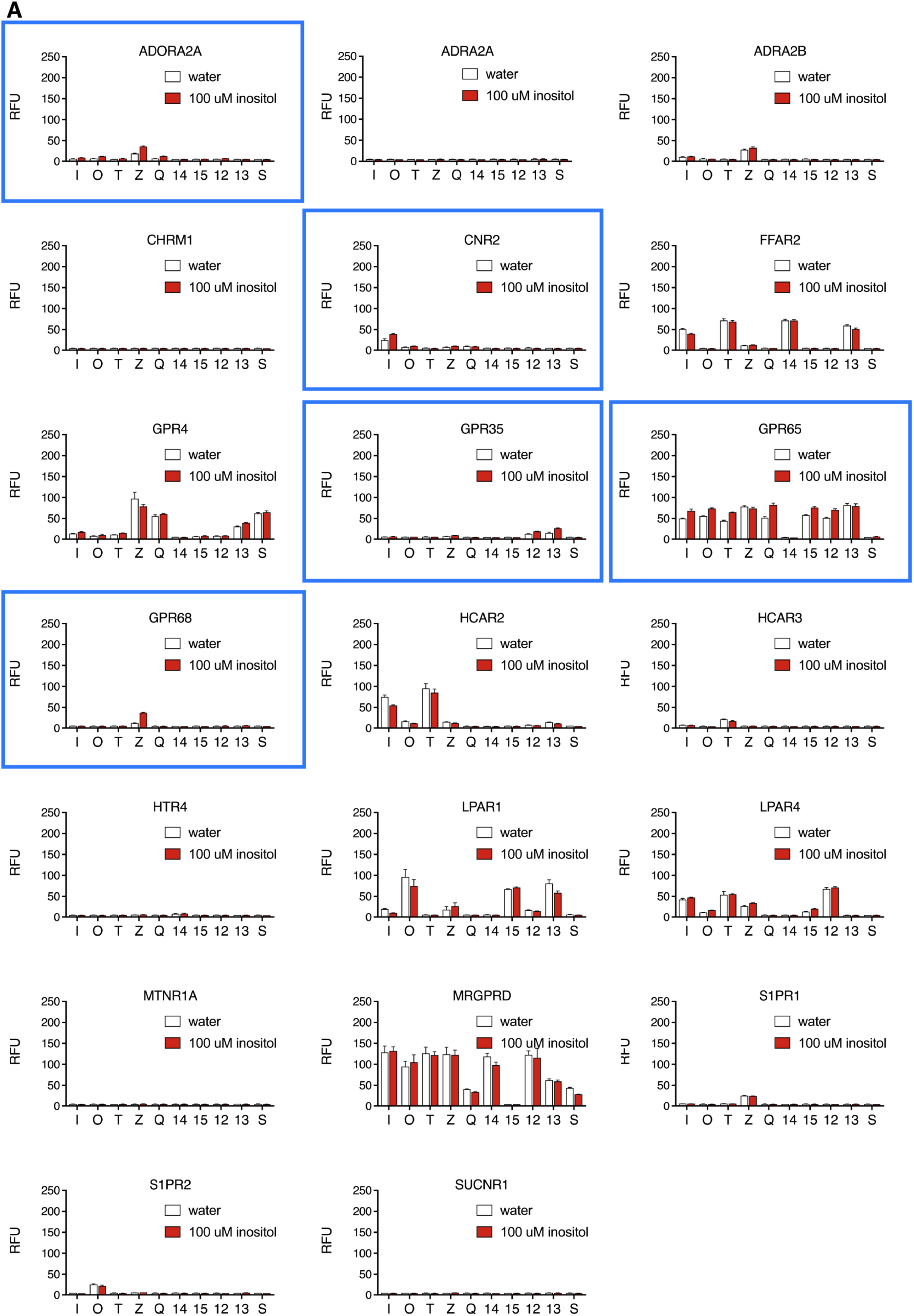
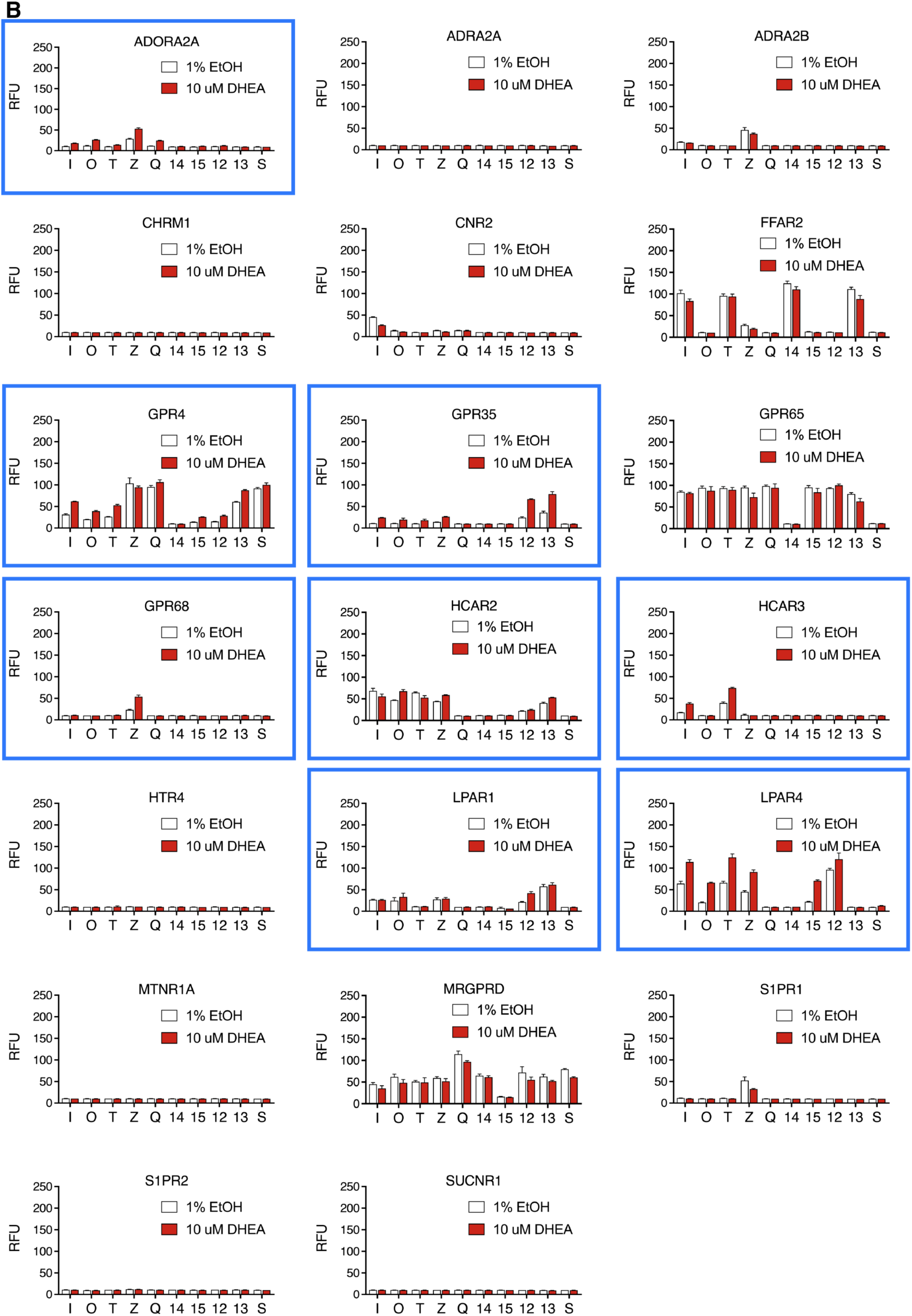
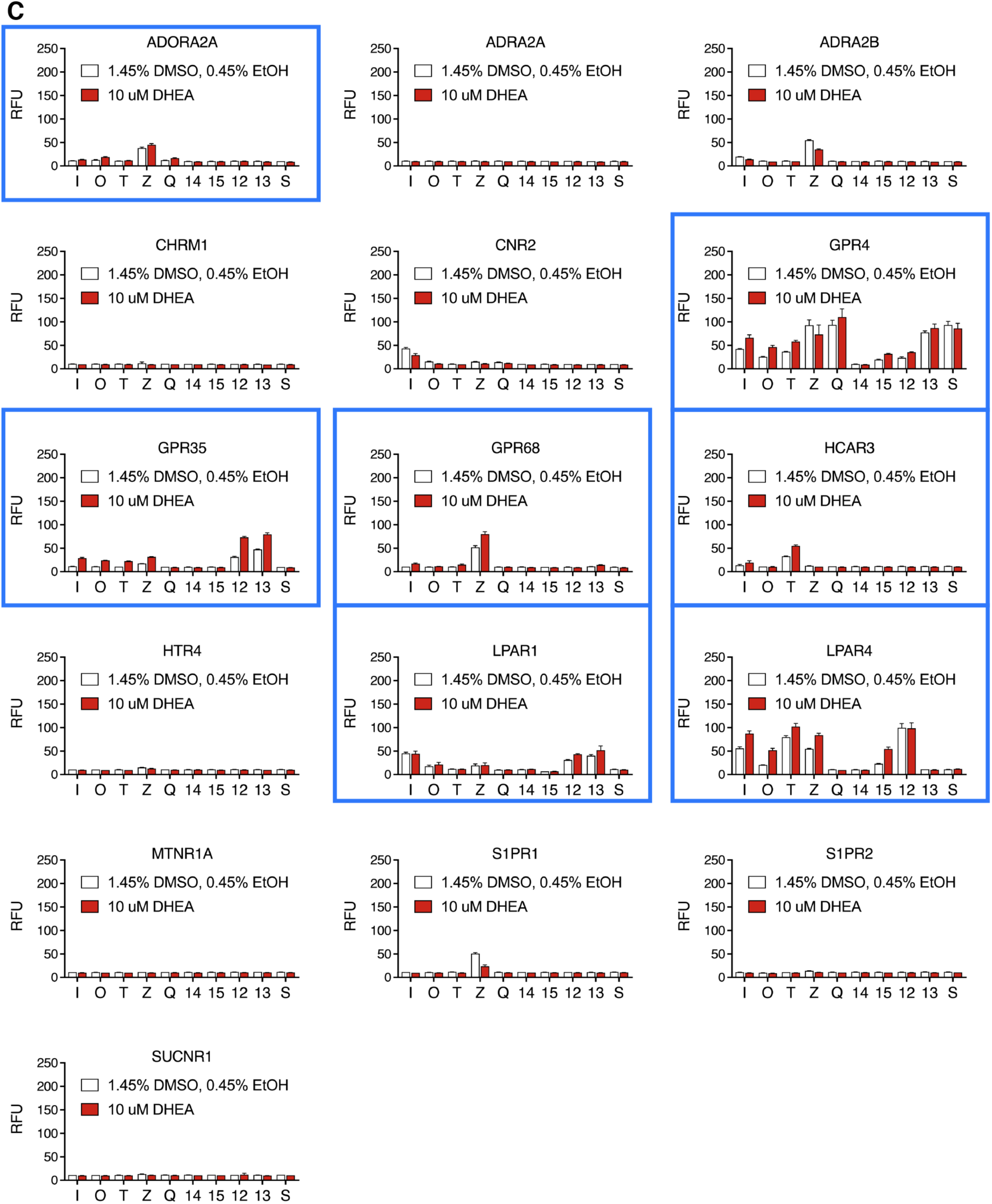
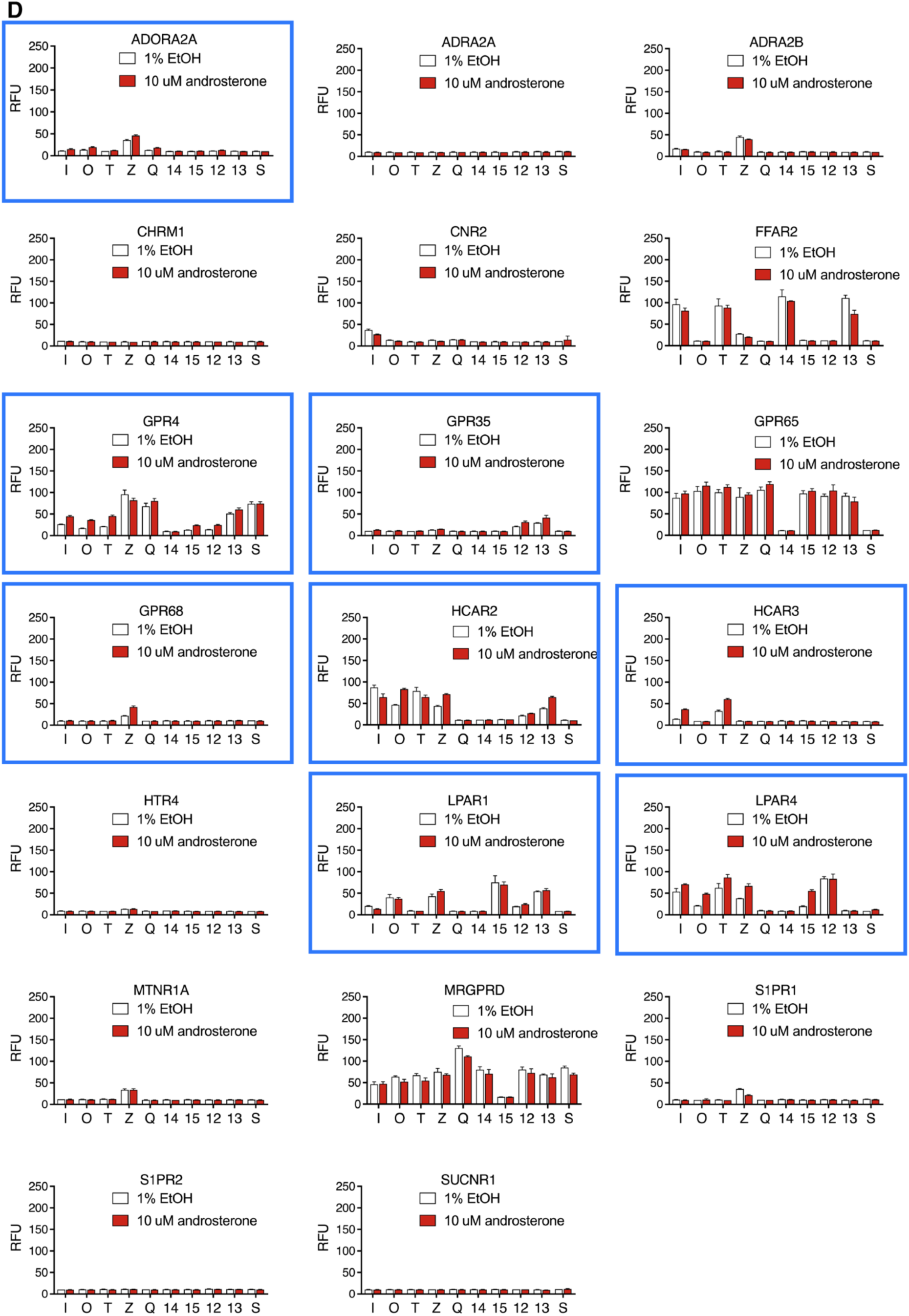
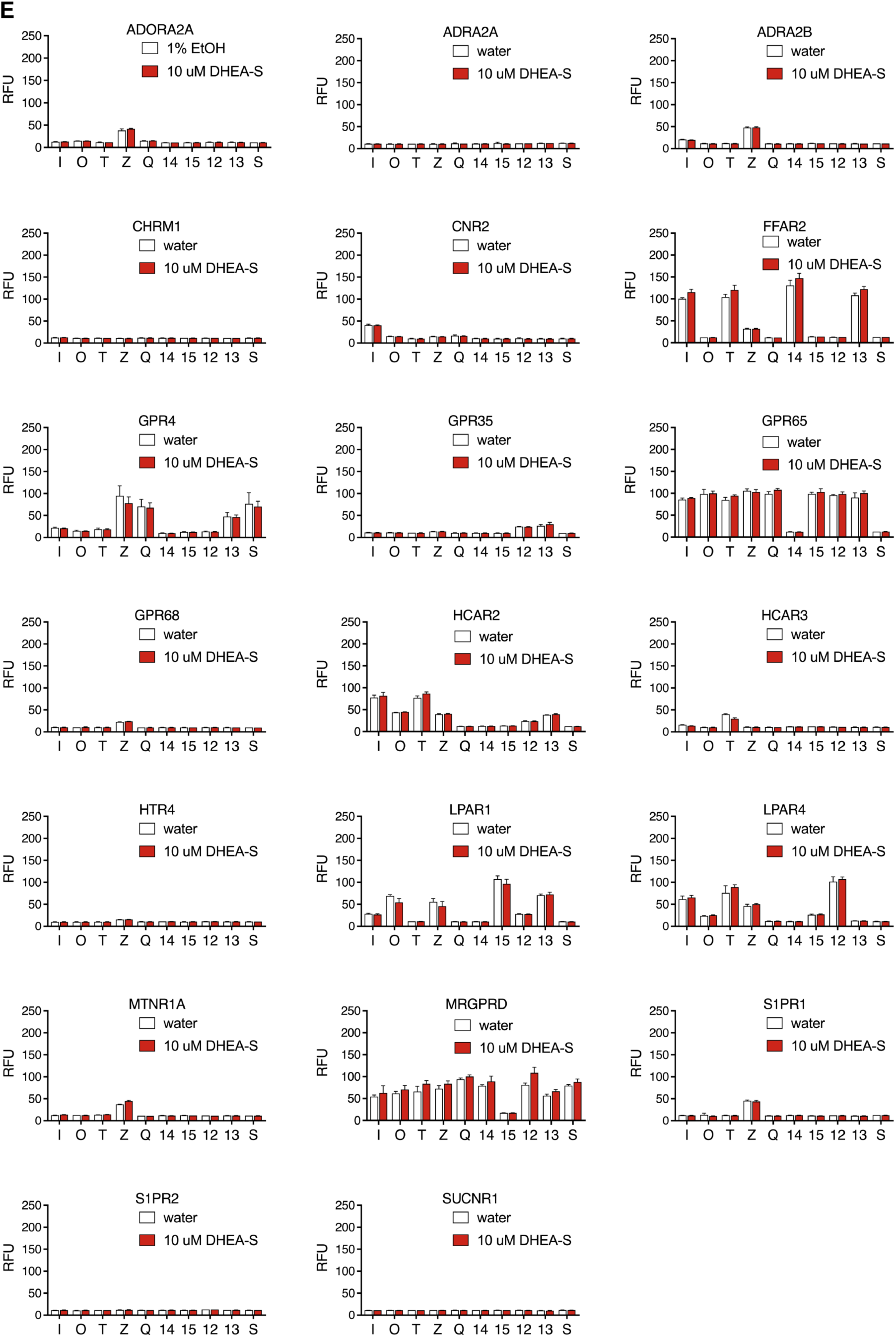
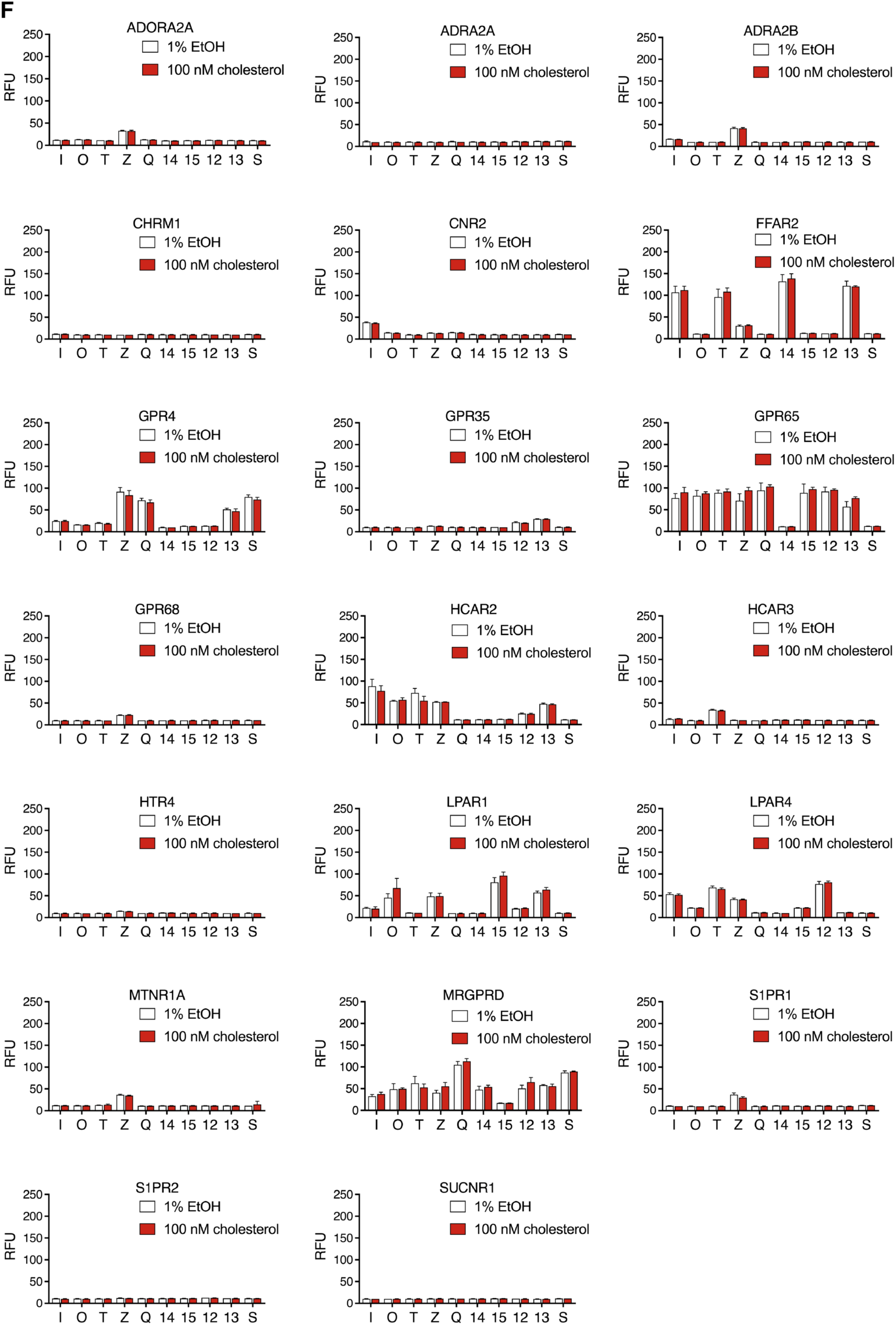
Control DCyFIRscreen profiles for new metabolite allosteric modulators. (A) DCyFIRscreen profiles for GPCR-Gα reporter strains treated with inositol. Error bars represent SD of n=4 technical replicates. Blue boxes indicate PAM interactions. (B) DCyFIRscreen profiles for GPCR-Gα reporter strains treated with DHEA in EtOH. Error bars represent SD of n=4 technical replicates. Blue boxes indicate PAM interactions. (C) DCyFIRscreen profiles for GPCR-Gα reporter strains treated with DHEA in DMSO/EtOH. Error bars represent SD of n=4 technical replicates. Blue boxes indicate PAM interactions. (D) DCyFIRscreen profiles for GPCR-Gα reporter strains treated with androsterone. Error bars represent SD of n=4 technical replicates. Blue boxes indicate PAM interactions. (E) DCyFIRscreen profiles for GPCR-Gα reporter strains treated with DHEA-S. Error bars represent SD of n=4 technical replicates. (F) DCyFIRscreen profiles for GPCR-Gα reporter strains treated with cholesterol. Error bars represent SD of n=4 technical replicates.

**Fig. S6.**
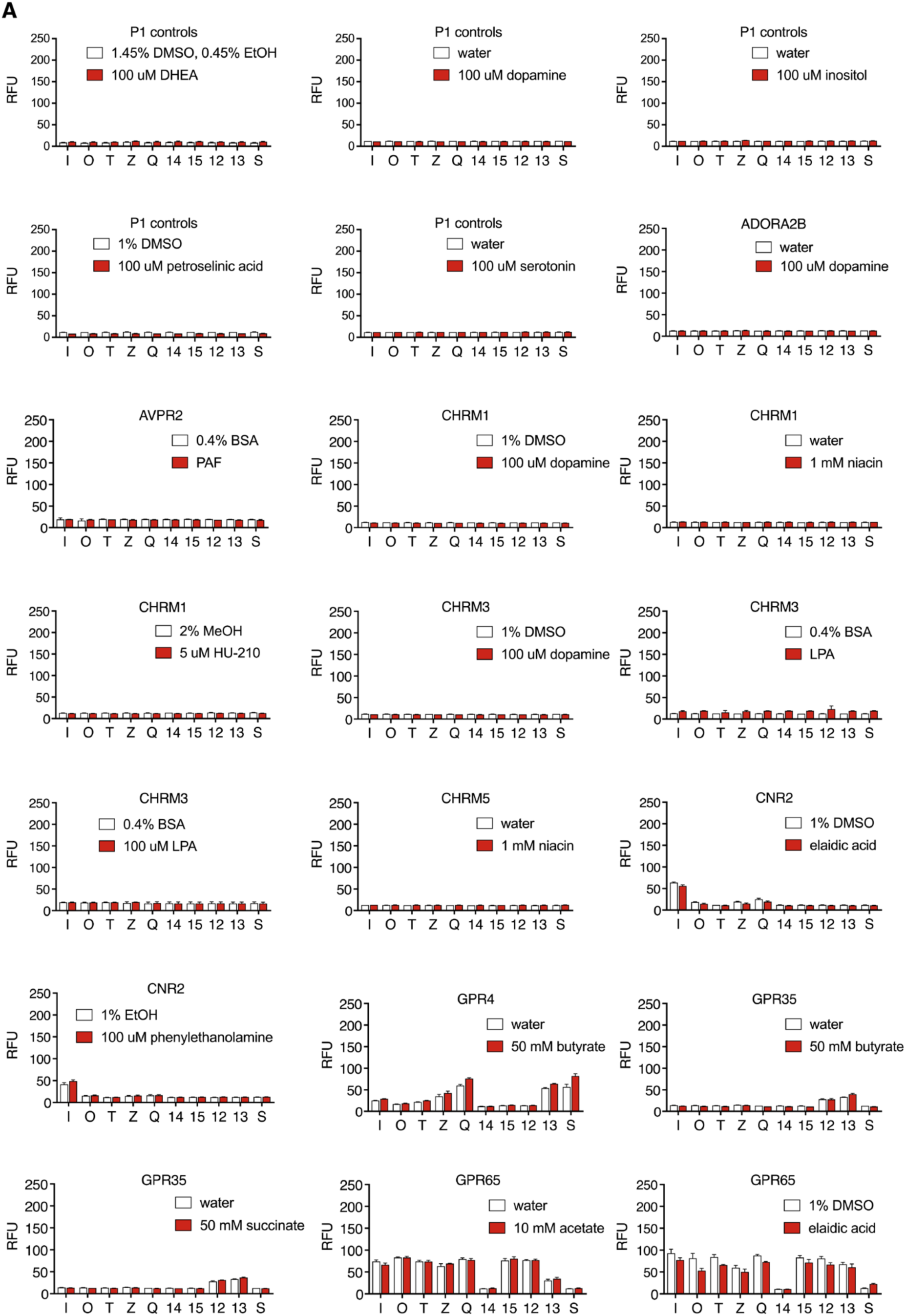
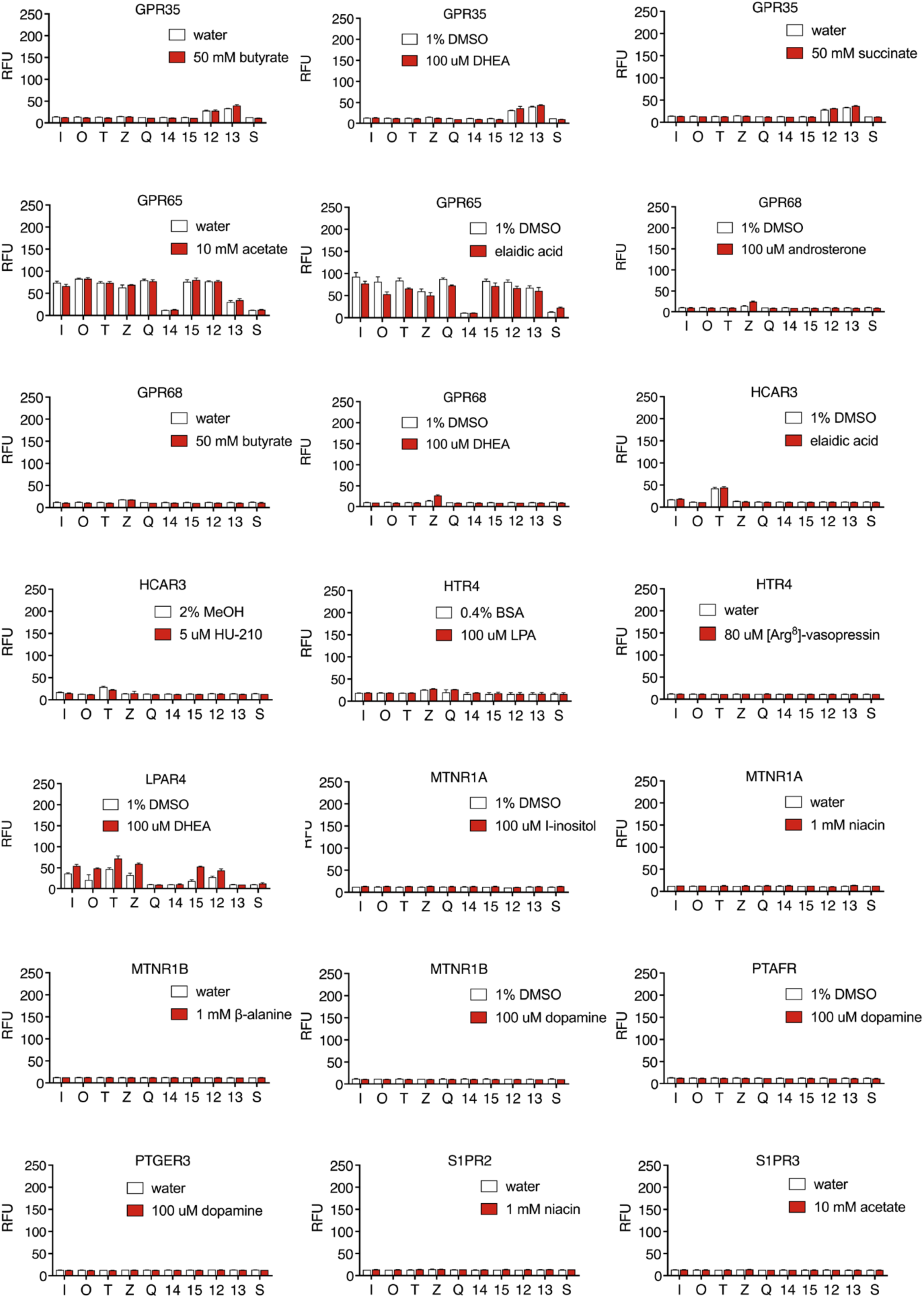
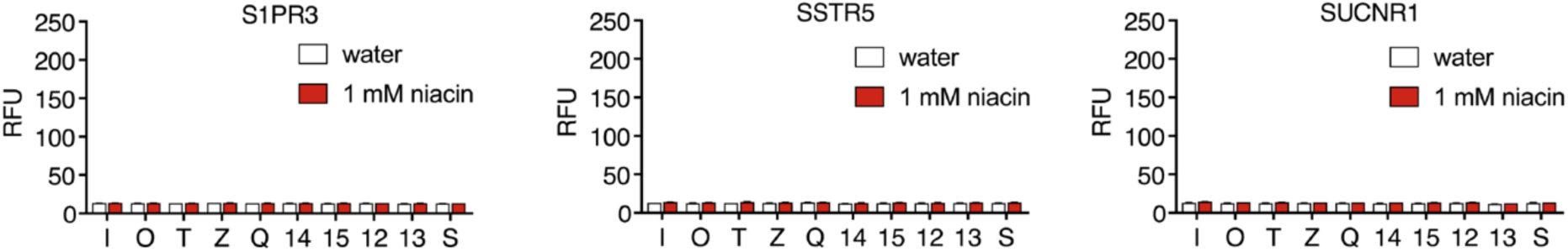
Control DCyFIRscreen profiles to assess DCyFIRplex sensitivity. (A) Control DCyFIRscreen profiles for receptors with low ΔCq values from DCyFIRplex profiling. Error bars represent SD of n=4 technical replicates.

**Table S1.** Media Formulations.

| **Medium** | **Concentration** | **Identifier** |
| --- | --- | --- |
| *Synthetic Complete Dextrose / Synthetic Complete Dextrose - Uracil Medium* |  | SCD/SCD -U |
| Ammonium sulfate | 5.00 g/L |  |
| CSM / CSM-URA supplement mixture | 0.79 / 0.77 g/L |  |
| Glucose | 20.00 g/L |  |
| Sodium hydroxide | 0.1 g/L |  |
| Yeast nitrogen base without amino acids & ammonium sulfate | 1.70 g/L |  |
| *Synthetic Complete Dextrose Low Fluorescence Screening Medium* |  | SCD LoFo pH 7.0 |
| Ammonium sulfate | 5.00 g/L |  |
| CSM supplement mixture | 0.79 g/L |  |
| Dextrose | 20.00 g/L |  |
| MES monohydrate | 9.76 g/L |  |
| Potassium phosphate dibasic | 8.72 g/L |  |
| Yeast nitrogen base- Low fluorescence without amino acids, folic acid, and riboflavin | 1.70 g/L |  |
| Potassium hydroxide solution | pH adjustment to 7.0 |  |
| Hydrochloric acid solution | pH adjustment to 7.0 |  |
| *Yeast Peptone Dextrose Medium* |  | YPD |
| Yeast extract | 10.00 g/L |  |
| Peptone | 20.00 g/L |  |
| D-(+)-Glucose | 10.00 g/L |  |
| *5FOA Counterselection Medium* |  | 5FOA |
| Yeast nitrogen base | 6.7 g/L |  |
| CSM supplement mixture | 0.79 g/L (has 20 mg uracil) |  |
| Uracil | 30 mg/L (50 mg total) |  |
| 5FOA | 1 g/L |  |
| Dextrose | 20 g/L |  |
| Bacto Agar | 15 g/L |  |

**Table S2.**
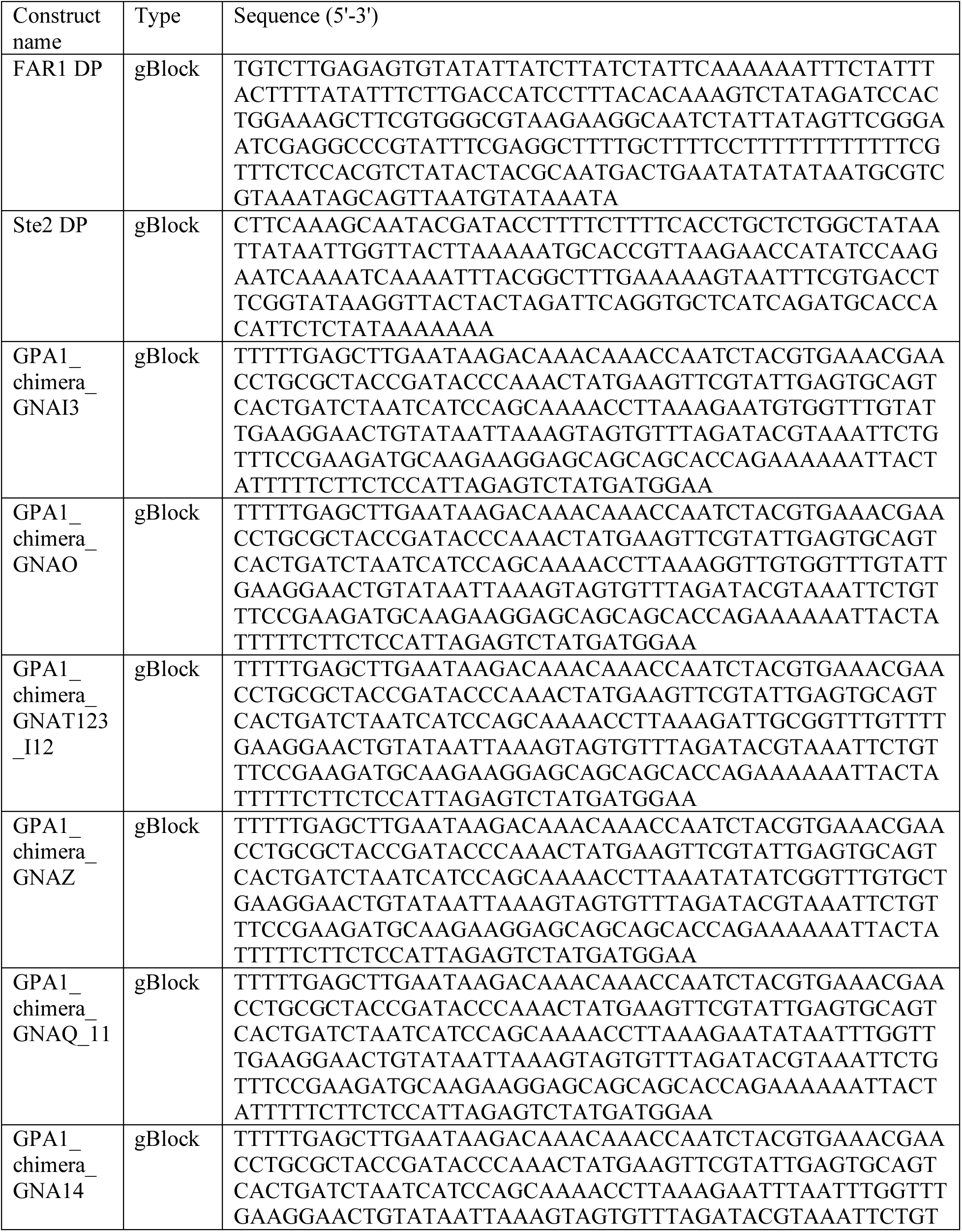
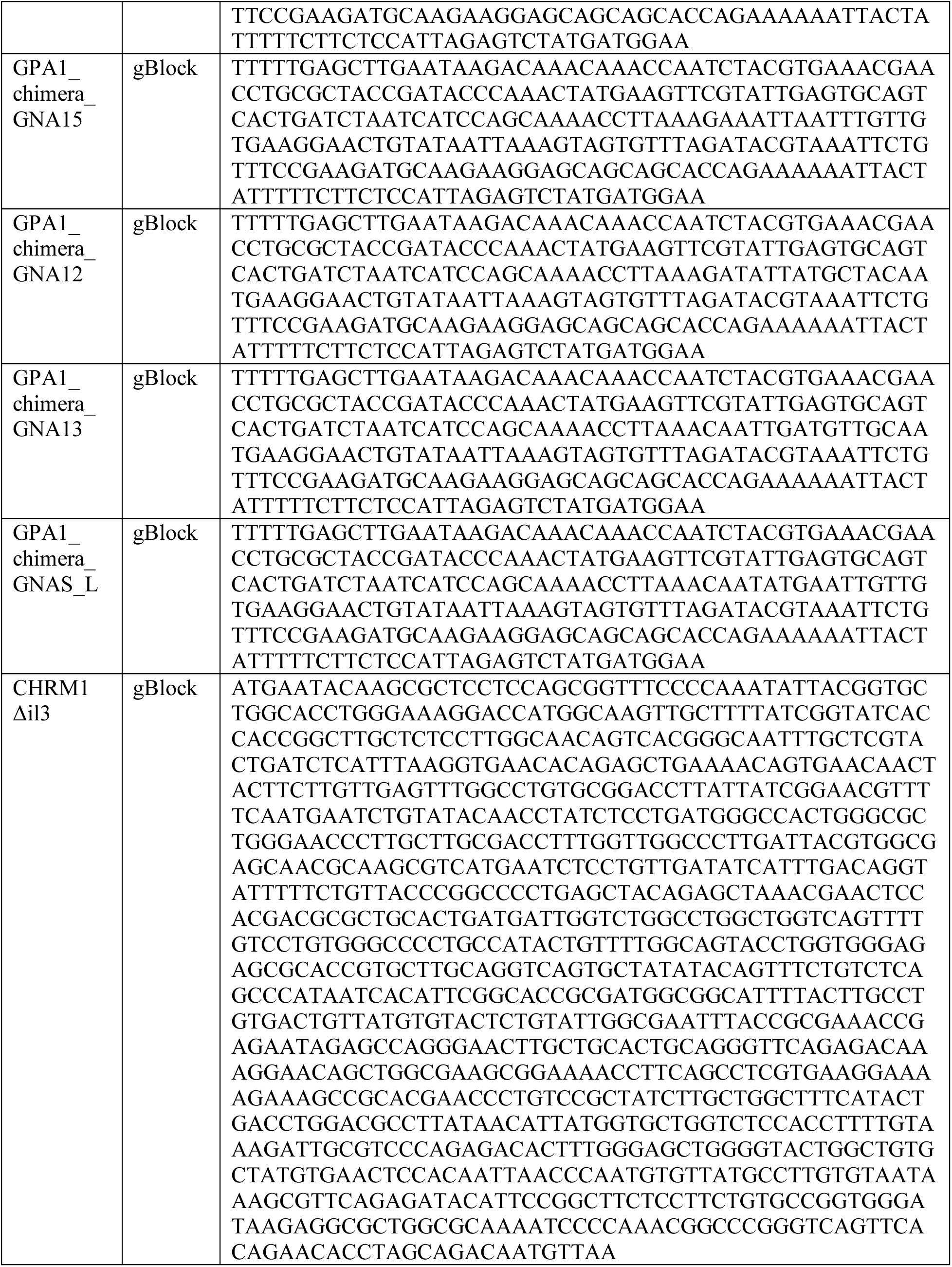
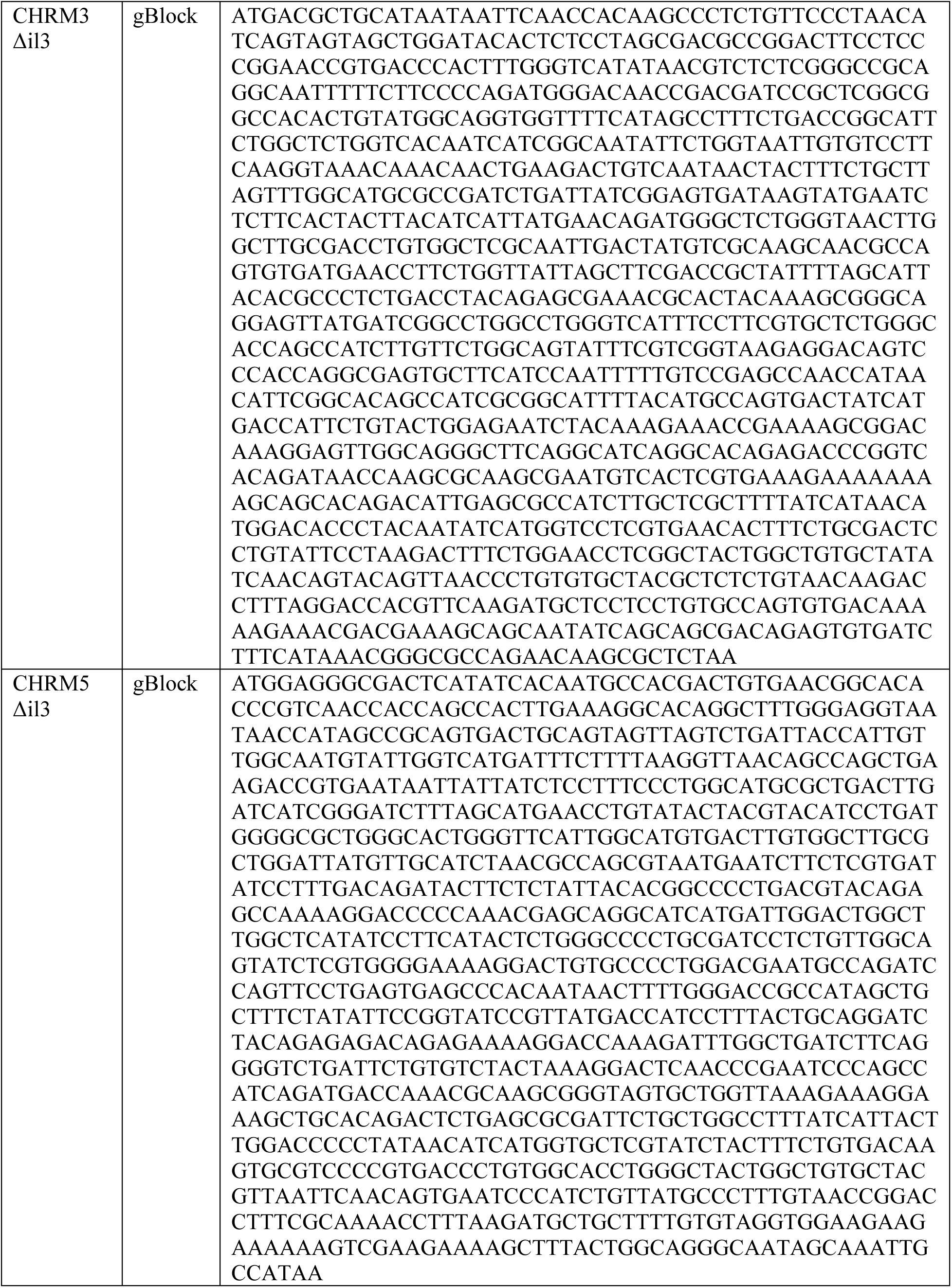
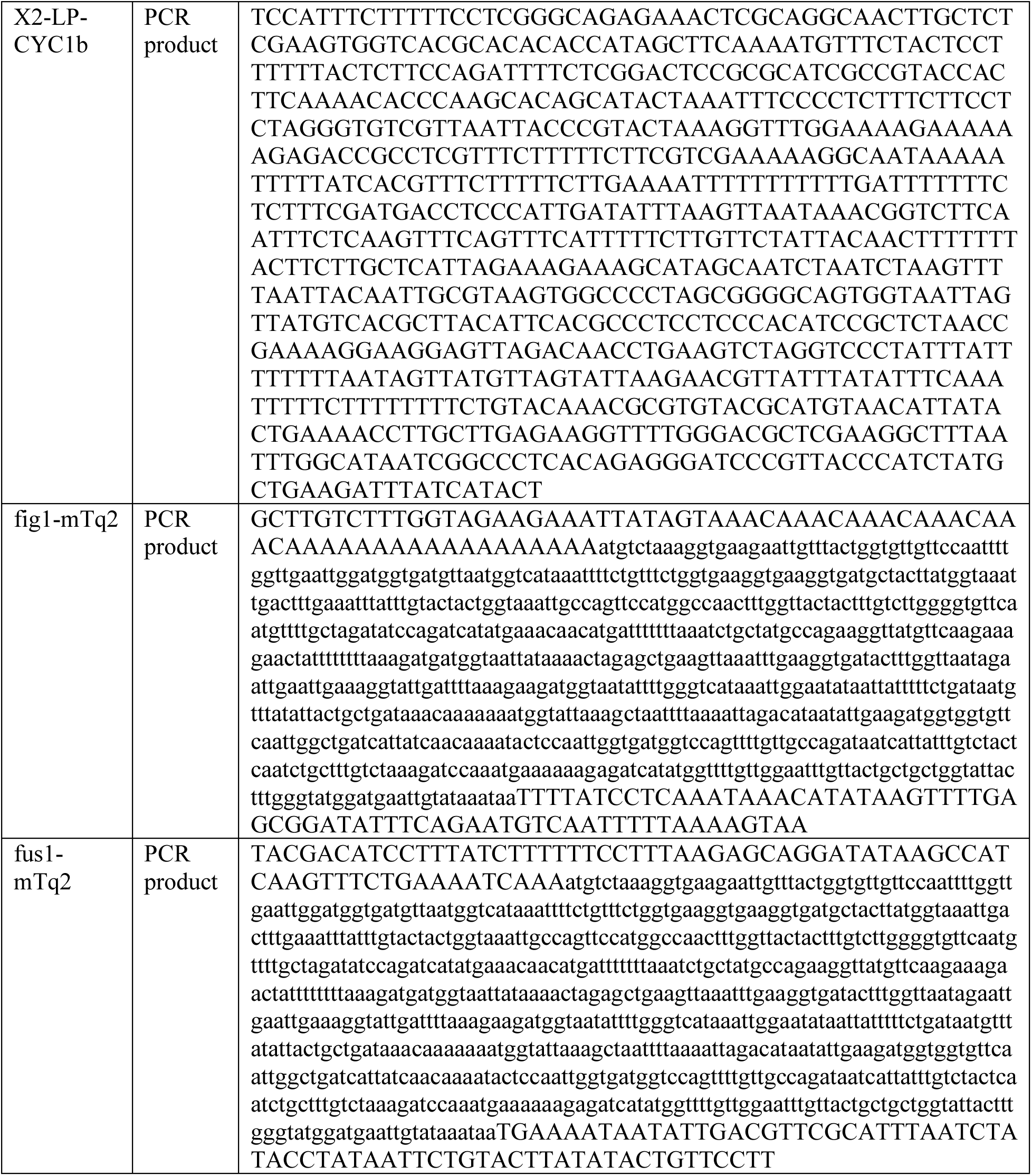
Synthetic DNA constructs used in this study.

**Dataset S1 (separate file).** Comprehensive strain list.

**Dataset S2 (separate file).** Human GPCR gene sequences.

**Dataset S3 (separate file).** Comprehensive plasmid list.

**Dataset S4 (separate file).** Oligonucleotides used in this study.

